# *Toxoplasma* GRA8 engages the host ESCRT accessory protein ALG-2 and is necessary for parasite metabolic integrity

**DOI:** 10.64898/2026.07.20.739547

**Authors:** Hargobinder Kaur, Rebekah B. Guevara, Yolanda Rivera-Cuevas, Einar B. Olafsson, Joshua Mayoral, Leonardo Augusto, Alfredo J. Guerra, Romir Patel, Kevin P. Bohannon, Jonathan Z. Sexton, Phyllis I. Hanson, Louis M. Weiss, Vern B. Carruthers

**Affiliations:** Department of Microbiology and Immunology, University of Michigan Medical School, Ann Arbor, Michigan, United States of America.; Department of Pathology, Albert Einstein College of Medicine, Bronx, New York, New York, United States of America.; Department of Pathology, and Microbiology, and Immunology, University of Nebraska Medical Center, Omaha, NE, USA.; Department of Biological Chemistry, University of Michigan Medical School, Ann Arbor, Michigan, United States of America.; Department of Medicinal Chemistry, College of Pharmacy, Ann Arbor, Michigan, United States of America.; Department of Medicine, Albert Einstein College of Medicine, Bronx, New York, New York, United States of America.

## Abstract

The endosomal sorting complex required for transport (ESCRT) is a hetero-multimeric membrane-remodeling machinery essential for endosomal sorting, intraluminal vesicle formation, cytokinetic abscission, and membrane repair. ESCRT is also hijacked by some pathogens including the protozoan *Toxoplasma gondii*, which subverts it at the parasitophorous vacuole membrane to support parasite ingestion of host cytosolic proteins. Although the ESCRT accessory protein, ALG-2 is recruited to the parasitophorous vacuole, nothing was known about how this happens. Herein we identify the dense granule protein TgGRA8 as a key effector that recruits ALG-2 and ALIX to the parasitophorous vacuole. We show that TgGRA8 directly binds ALG-2 via conserved ALG-2-binding elements like those found in other ALG-2 interacting proteins including ALIX and SEC31A. Biochemical assays show high-affinity, Ca²⁺-dependent TgGRA8-ALG-2 binding, and structural modeling suggests TgGRA8 may assemble multivalently to coordinate multiple ALG-2 dimers, stabilizing ALG-2/ALIX recruitment through a non-canonical bridging mechanism. Conservation of these motifs among tissue cyst-forming coccidians implies a lineage-linked adaptation that supports infection by these parasites. Metabolomics further indicates that TgGRA8 loss disrupts amino acid, purine, and central carbon metabolism, like those seen in other ingestion deficient mutants. Together, these findings uncover a conserved, multivalent strategy by which *Toxoplasma* engages host ALG-2 to organize ESCRT at the parasitophorous vacuole, thereby coupling nutrient acquisition to metabolic fitness and exposing a novel agent for probing ESCRT biology.

## Introduction

The endosomal sorting complex required for transport (ESCRT) is an evolutionary ancient, hetero-multimeric membrane-remodeling machinery. Comprised of ESCRT-0, -I, -II, and -III, the AAA+ ATPase Vps4, and associated factors, ESCRT drives membrane-neck scission in diverse cellular contexts, including multivesicular body biogenesis, HIV budding, cytokinesis, plasma and lysosomal membrane repair (1). The ESCRT machinery works together with several accessory proteins including apoptosis-linked gene 2 (ALG-2, also known as PDCD6).

ALG-2 is an EF-hand, Ca²⁺-binding protein that homodimerizes to couple intracellular signals to ESCRT function. This coupling occurs through Ca²⁺-dependent binding of ALG-2 to conserved proline-rich motifs in partner proteins including ALG-2-interacting protein X (ALIX, also known as PDCD6IP and AIP1), the first identified Ca²⁺-dependent ALG-2 interactor (2,3). Site mapping and mutational analyses delineate three proline-based ALG-2-binding motifs (ABMs) in ALG-2 interacting proteins (4). The ABM-1, consisting of the proximal motifs PPYP and YP (in e.g., ALIX), binds respectively to adjacent sites in Pocket 1 and Pocket 2 of ALG-2. The ABM-2 motif PPPPGF (in e.g., Sec31A) targets a distinct Pocket-3 of ALG-2. The ABM-3 consists of MP tandem repeats (in a polymorphic isoform of IST1) but such motifs alone are insufficient for binding to ALG-2 (5). In ALIX, a ABM-1 motif immediately followed by a ABM-3 like segment may potentiate ALG-2 engagement, but this has not been directly tested (5). An alternatively spliced isoform lacking residues 121-122 (ALG-2^ΔGF122^) has a partially occluded Pocket-1 that abolishes ALIX binding without affecting binding to pocket 3 (6–8). ALG-2 localizes to the cytoplasm and nucleus where it functions to augment interactions between membrane-proximal scaffolds (9–11). For example, whereas the direct interaction between ALIX (PSAP motif) and TSG101 (UEV domain) is weak, Ca²⁺-activated dimeric ALG-2 binds to ABMs on both ALIX and TSG101 (a component of ESCRT-I), thereby serving as an adapter to stabilize the complex (11–13).

Diverse pathogens hijack host ESCRT to enable their replication and dissemination (14). For example, during viral assembly many enveloped viruses recruit ESCRT via short “late-domain” motifs (PTAP, PPXY, YPXnL) that mimic host sequences to engage TSG101 and/or ALIX, thereby driving budding away from the cytosol (15–18). As it assembles at the plasma membrane, HIV-1 Gag nucleates ESCRT recruitment through p6 late domain-encoded PTAP-TSG101 and YPXnL-ALIX interactions, with downstream ESCRT-III and VPS4 activities required for budding (15,19–21). In addition to enveloped viruses, intracellular bacteria exploit the ESCRT machinery to maintain pathogen-containing vacuoles (PCVs) in *Salmonella* (22,23) or antagonize ESCRT to evade clearance in *Mycobacterium* (24,25), underscoring context-dependent roles. The fungal pathogen *Candida albicans* activates ALG-2/ESCRT-III dependent membrane shedding and lysosome-mediated repair, preserving epithelial barrier integrity during both commensal colonization and infection (26). Among parasites, *Leishmania donovani* engages ESCRT during macrophage infection via a non-canonical, ALIX-initiated pathway where ALIX is involved in catalyzing the scission of lysosome-derived parasitophorous vacuoles (PVs) (27). However, the underlying mechanism of this engagement is unknown.

Emerging evidence indicates that the protozoan pathogen *Toxoplasma gondii* (*T. gondii*) also exploits host ESCRT machinery at its PV during infection (28). *T. gondii*, which can infect any nucleated cell, recruits to its PV key components of ESCRT-I (TSG101), ESCRT-III (CHMP4B), accessory proteins ALG-2 and ALIX, and VPS4 (28–30). Notably, the dense granule protein TgGRA14 uses a PTAP motif to recruit TSG101 to the PV membrane (PVM), facilitating the budding of vesicles containing host cytosolic material into the PV lumen (29). Some of these TgGRA14- and ESCRT-dependent vesicles are endocytosed by the parasite via the so-called ingestion pathway to provide certain metabolites (31,32), whereas others containing host Rab11a positive vesicles remain in the PV as a source of lipids for the parasite (30). Another dense granule protein, TgEAF1, was recently identified as contributing to the recruitment of TSG101 to the PVM (33). A third dense granule protein, TgGRA64, also binds to host ESCRT proteins (34), but its role in recruiting ESCRT machinery to the PV is unknown. Likewise, although host ALG-2 and ALIX are recruited to the PV (28,29), the effector protein(s) responsible for such recruitment is unknown.

To address this gap, herein we identify the *T. gondii* dense granule protein 8 (TgGRA8) (35) as the first microbial effector shown to directly engage ALG-2. TgGRA8 facilitates the association of ESCRT components at the PVM by serving as a principal recruiter of ALG-2 together with ALIX in both virulent type I (RH) and avirulent type II (ME49) *T. gondii* strains. Remarkably, TgGRA8 harbors three tandemly repeated elements comprising both ABM-1 and ABM-2 motifs, a combination that was previously unknown in ALG-2-binding proteins. Biochemical analyses revealed high-affinity binding of peptides from repeated elements to ALG-2. Consistent with a key role in ALG-2 recruitment, disrupting TgGRA8 resulted in pronounced perturbation of *T. gondii* metabolites including amino acids, nucleosides, and those comprising central carbon metabolism, mirroring the metabolic perturbations seen in ingestion pathway mutants (32).

## Results

### TgGRA14 and TgGRA64 are not responsible for ALG-2 recruitment to the *Toxoplasma* PV

To determine how *Toxoplasma* recruits ALG-2 to its PV, we began by looking at two previously identified ESCRT interacting parasite proteins, TgGRA14 and TgGRA64. To interrogate TgGRA14, we measured the intensity of ALG-2, ALIX, and TSG101 at the PVM in HeLa cells infected with type I RH (WT) or RH*Δgra14* (*Δgra14*) parasites. For a robust assessment, we used a semi-automated high content imaging and image analysis pipeline to measure the mean intensity ratio of ALG-2, ALIX, and GFP-TSG101 at PV relative to the cell cytoplasm, excluding the PV and nucleus. ALG-2 recruitment was assessed in HeLa cells transiently expressing ALG-2-HA, ALIX was detected by antibody staining, and TSG101 was tracked in HeLa cells expressing endogenously tagged GFP-TSG101 (36). Quantitative imaging showed that TgGRA14 plays a statistically significant but minimal role in the PV recruitment of ALG-2 and ALIX compared to its pronounced function in recruiting TSG101 (**Fig 1).** Analysis of well-level data confirmed the differences observed for the biological replicates (**S1A Fig)**. Using a synthetic peptide encoding the TgGRA14 PTAP motif, we also showed that it binds to the TSG101 UEV domain with a K_d_ of 18.2±0.8 µM, which is a moderately higher affinity than that of an HIV-1 Gag PTAP peptide (43.7±1.6 µM) tested in the same experiments (data not shown). Collectively, these data suggest that TgGRA14 binds directly to TSG101 and is the principal recruiter of TSG101 to the PV, but that it does not exclusively account for recruitment of ALG-2 or ALIX.

**Fig 1.**
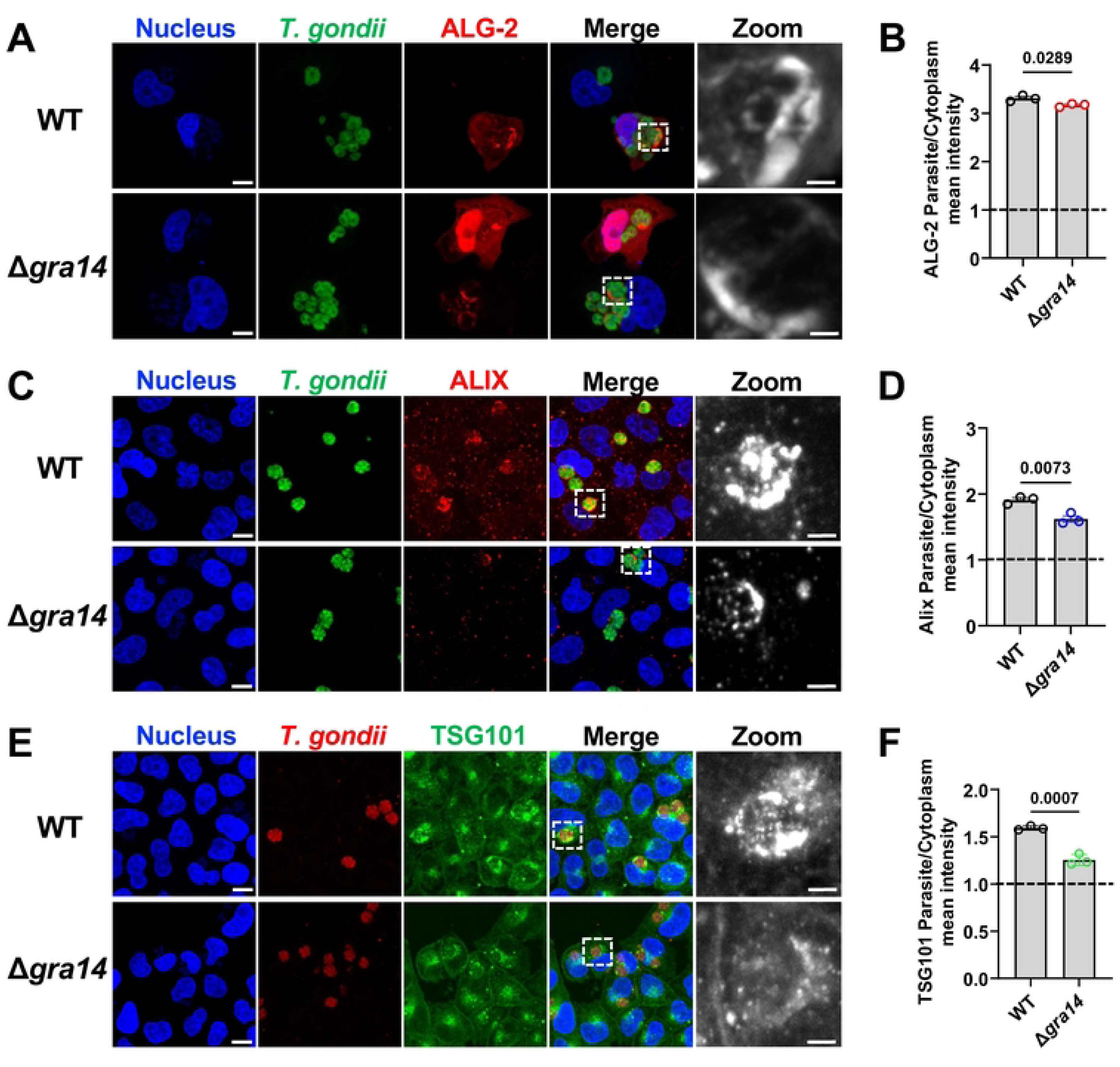
TgGRA14 is not responsible for ALG-2 recruitment to the PV. **(A, C, E).** ALG-2-HA transfected HeLa, WT-HeLa and GFP-TSG101-expressing HeLa cells were either infected with RH (WT) or RHΔ*gra14* parasites. Host cell nuclei were stained with DAPI, and parasites were visualized using anti-*T. gondii* Ab. ALG-2, and ALIX were probed using Anti-HA, and Anti-ALIX mAb (Monoclonal Ab), while TSG101 was endogenously tagged with GFP. Images are representative of three biological replicates analyzed by confocal microscopy. Scale bar, 20 µm. Black and white zoomed insets representing the ALG-2, ALIX and TSG101 recruitment on the PVM. Scale bar, 2 µm. **(B, D, F).** Quantification of recruitment of ALG-2, ALIX and TSG101 to the PVM. Data represents the mean ALG-2/ALIX/TSG101 intensity at the PV relative to the cytosol (PV/cytosol). Data is from 3 biological replicates each with >10,000 PVs analyzed. Each point represents the well average (∼16 wells per biological replicate) ± SEM. Statistical analysis was performed using unpaired students t-test. The dashed line at a ratio of 1 indicates little to no enrichment at the PV versus the host cytoplasm.

Although prior work suggested that TgGRA64 co-immunoprecipitated with several host ESCRT components and contains a poly basic putative ALIX binding motif (34), the role of TgGRA64 in recruiting ALIX or other ESCRT proteins to the PVM has not been tested. To determine whether TgGRA64, alone or together with TgGRA14, contributes to ESCRT protein recruitment, we quantified the relative intensities of ALG-2, ALIX, and TSG101 at the PVM using an approach described above. HeLa cells were infected with RHΔ*ku80*WT (WT), RHΔ*ku80*Δ*gra64* (*Δgra64*), or RHΔ*ku80*Δ*gra64*Δ*gra14* (Δ*gra64*Δ*gra14*) parasites. Deletion of TgGRA64 alone slightly increased PV recruitment of ALG-2 **(Fig 2A, B)**. This augmented recruitment was significantly but modestly decreased in Δ*gra64*Δ*gra14* parasites, confirming a minor role for TgGRA14 in recruiting ALG-2 to the PV. TgGRA64 deficient parasites recruited significantly less ALIX to the PV, a decrease that was further accentuated in Δ*gra64*Δ*gra14* parasites **(Fig 2C, D)**, suggesting roles for both TgGRA64 and TgGRA14 in the PV recruitment of ALIX. The abundance of TSG101 at the PVM was unchanged in TgGRA64 deficient parasites but was substantially reduced in parasites lacking both TgGRA64 and TgGRA14 **(Fig 2E, F)**, consistent with the known interaction of TgGRA14 with TSG101. Compilation of well-level data confirmed the differences observed for the biological replicates (**S1B Fig).** Together, these findings indicate that TgGRA64 and TgGRA14 additively contribute to ALIX recruitment to the PV, but TgGRA64 does not influence the PV recruitment of ALG-2 or TSG101.

**Fig 2.**
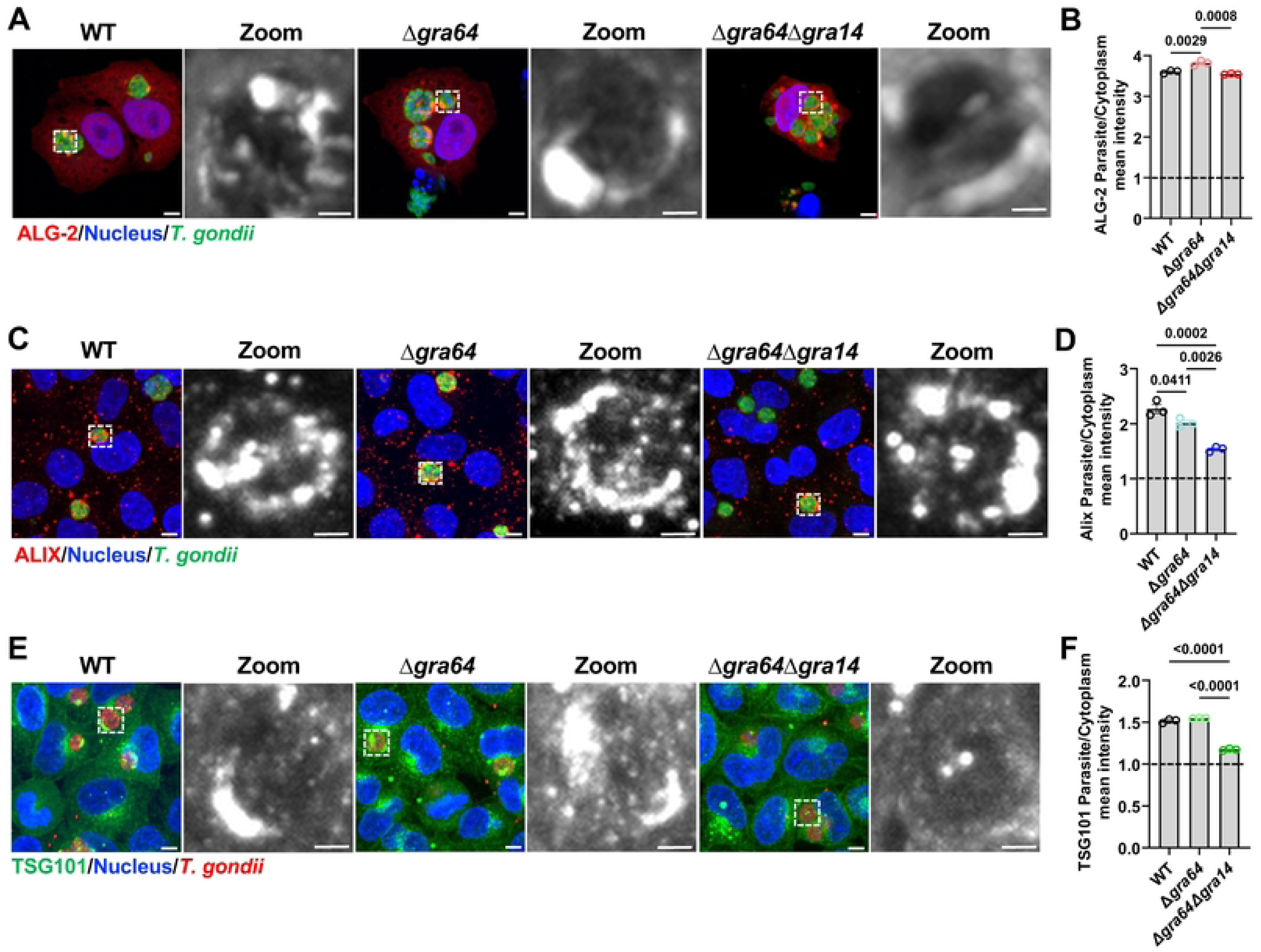
GRA64 is not responsible for ALG-2 recruitment to the PV. **(A, C, E).** ALG-2-HA transfected, WT, and GFP-TSG101-expressing HeLa cells were infected with either RH*Δku80*WT, RHΔ*ku80*Δ*gra64* or RHΔ*ku80*Δ*gra64*Δ*gra14* parasites. Host cell nuclei were stained with DAPI, and parasites were visualized using anti-*T. gondii* antibody. ALG-2 and ALIX are probed using anti-HA and anti-ALIX antibodies, and TSG101 was endogenous GFP tagged. Images shown are representative of 3 independent biological replicates analyzed by confocal microscopy. Scale bar, 20 µm. Black and white zoomed insets representing the ALG-2, ALIX and TSG101 recruitment on the PVM. Scale bar, 2 µm. (**B, D, F).** Recruitment of ALG-2, ALIX, and TSG101 to the PV was quantified as the mean fluorescence ratio of PV-to-cytosol across 3 biological replicates (>10,000 PVs each). Each point represents the well average (∼16 wells per biological replicates) ± SEM. Statistical comparisons were performed using one-way ANOVA with Tukey’s multiple comparisons test. The dashed line at a ratio of 1 indicates little to no enrichment at the PV versus the host cytoplasm.

### Identification of TgGRA8 as an ALG-2-interacting protein

To identify parasite proteins that interact with ALG-2, we immunoprecipitated ALG-2 from human foreskin fibroblast (HFF) cells infected with type II Pru strain parasites. Quantitative proteomic analysis showed the expected enrichment of ALG-2, along with several other host proteins including the ESCRT-III protein CHMP4B and the ESCRT-I proteins VPS37C and TSG101, which cooperate with ALG-2 in diverse cellular membrane remodeling processes **(Fig 3A, S1 Data)**. The analysis also identified three *T. gondii* proteins: TgGRA8, TgGRA12, and TgGRA15 **(Fig 3B)**. We focused on TgGRA8 because it was the most highly enriched parasite protein associated with ALG-2. TgGRA8 is a protein of unknown function that spans the PVM (35), thereby giving it access to the host cytosol. An interaction between ALG-2 and TgGRA8 was confirmed by immunoprecipitation and immunoblotting, with the unrelated dense granule protein TgGRA1 serving as a negative control **(Fig 3C)**. Reciprocal quantitative proteomic analysis of TgGRA8 immunoprecipitated from HFF cells infected with parasites expressing endogenously HA tagged TgGRA8 (Pru::HA-GRA8) showed significant enrichment of ALG-2 and ALIX along with proteins from ESCRT-I (TSG101, VPS37A/B/C, VPS28, UBAP1, MVB12A, and UMAD1), ESCRT-II (VPS25), and ESCRT-III (CHMP1A, CHMP2A/B, CHMP3, CHMP4B, and IST1) complexes **(Fig 3D,G; S2 Data)**. Among *T. gondii* proteins, TgGRA8 was highly enriched along with TgGRA64, TgGRA14, and other dense granule proteins **(Fig 3E)**, indicating potential cooperativity among GRA proteins targeting the host ESCRT machinery. Interaction of TgGRA8 with ALG-2 and TgGRA14 was confirmed by immunoprecipitation and immunoblotting **(Fig 3F).** Together, these findings demonstrate that TgGRA8 interacts with ALG-2 and several other ESCRT and ESCRT-associated proteins during infection.

**Fig 3.**
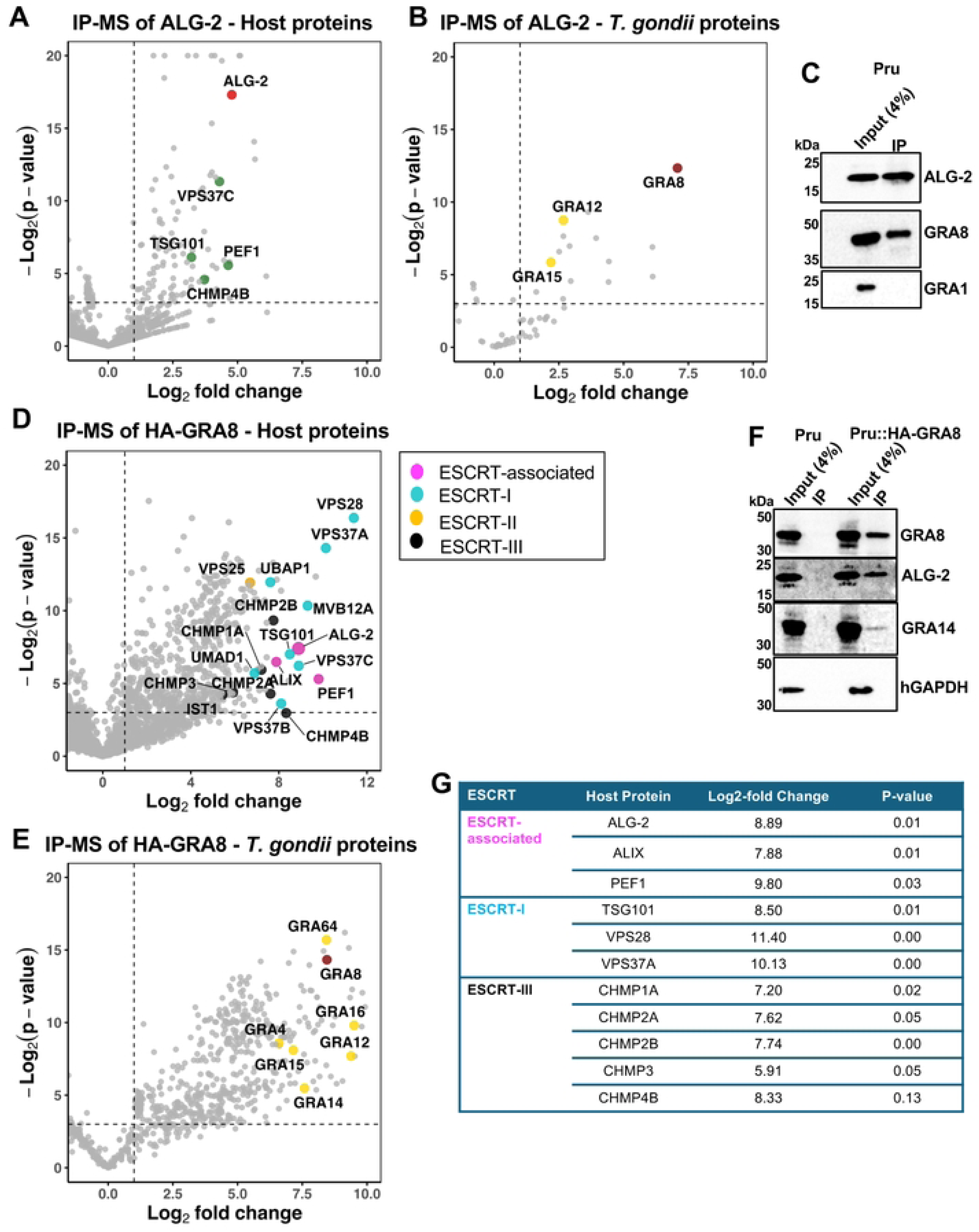
ALG-2 IP reveals TgGRA8 as a potential *Toxoplasma* effector protein. **(A, B)** Immunoprecipitation of ALG-2 was performed on PruΔ*ku80*WT infected human foreskin fibroblast (HFF) cells and further Mass spectrometry (MS) was performed. Volcano plots showing host **(A)** and *T. gondii* **(B)** proteins identified with IP-MS/MS. **(C)** Immunoblot analysis of ALG-2 immunoprecipitates confirming ALG-2 interaction with TgGRA8. GRA1 served as a negative control. **(D, E)** IP of HA-GRA8 with anti-HA was performed on Pru::HA-GRA8 infected HFF cells expressing HA endotagged TgGRA8 followed by MS analysis. Volcano plots demonstrating host **(D)** and *T. gondii* **(E)** proteins identified by IP-MS of HA-tagged TgGRA8. TgGRA8 interacts with multiple ESCRT-associated proteins (magenta), including ALG-2, ALIX, and several components of the ESCRT-I (cyan), ESCRT-II (Yellow), and ESCRT-III complexes (Black). Colored dots represent proteins with >1 log2 fold change in tagged vs control lysates and a negative log2 p-value >3.32 (equivalent to p<0.1) **(F)** Immunoblots confirming TgGRA8 interaction with host ALG-2 and TgGRA14, hGAPDH served as a negative control. **(G)** Summary table of host ESCRT-associated proteins identified by IP-MS of TgGRA8, showing log2 fold change and p-values.

### TgGRA8 is the main recruiter of ALG-2 and ALIX to the PV

To define the role of TgGRA8 in recruiting ALG-2, ALIX, and TSG101 to the PV, we analyzed recruitment in HeLa cells infected with RHΔ*ku80* (WT), RHΔ*ku80*Δ*gra8* (Δgra8 knockout), and RHΔ*ku80*Δ*gra8GRA8* (Δ*gra8GRA8* complement) parasites. Deletion of TgGRA8 markedly reduced the PV recruitment of transiently overexpressed ALG-2-HA, which was restored by its re-expression in Δ*gra8GRA8* parasites **(Fig 4A, B)**. PV recruitment of ALIX was likewise significantly diminished in *Δgra8* parasites and rescued in Δ*gra8*GRA8 parasites **(Fig 4C, D)**. TgGRA8 deficient parasites showed a trend toward reduced PV recruitment of TSG101, but the difference was not statistically significant **(Fig 4E, F)**. To corroborate these findings, we generated the corresponding TgGRA8 transgenic parasites in a type II ME49 strain background. ME49*Δgra8* parasites showed significantly less ALG-2, ALIX, and TSG101 at the PV, which was rescued by re-expression of TgGRA8 (**S2 Fig**). Analysis of well-level data confirmed the differences observed for biological replicates in both strain backgrounds (**S3A, B Fig).** Consistent with the findings for transfected ALG-2-HA, an optimized staining protocol for endogenous ALG-2 showed a similar loss of ALG-2 recruitment to the PV in Δ*gra8* parasites in both type I and type II backgrounds (S4A, B Fig). Collectively, these findings indicate that TgGRA8 plays a principal role in recruiting ALG-2 to the PVM, while also contributing to the recruitment of ALIX and TSG101.

**Fig 4.**
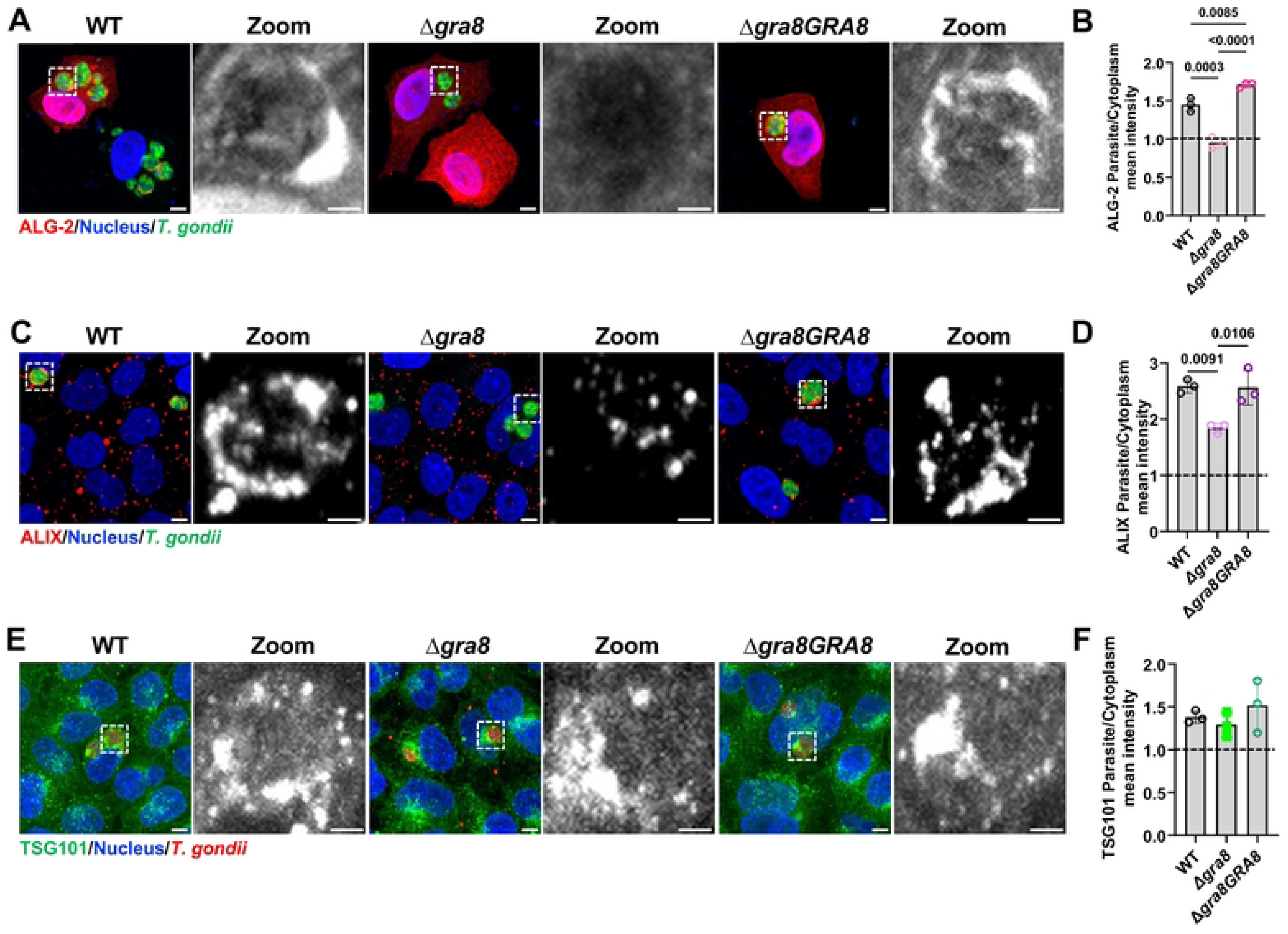
TgGRA8 is essential for PV recruitment of ALG-2. **(A, C, E)** HeLa cells expressing ALG-2-HA, wild-type (WT) HeLa cells, or GFP-TSG101-expressing HeLa cells were infected with RHΔ*ku80*WT, RHΔ*ku80*Δgra8, or RHΔ*ku80*Δ*gra8*::*GRA8.* ALG-2 and ALIX were detected using anti-HA and anti-ALIX antibodies, respectively, while TSG101 was visualized via endogenous GFP tagging. Representative images from 3 independent biological replicates are shown. Scale bars, 20 µm. Black and white zoomed insets representing the ALG-2, ALIX and TSG101 recruitment on the PV. Scale bar, 2 µm. **(B, D, F)** Quantification of ALG-2, ALIX, and TSG101 recruitment to the PV, represented as parasite-to-cytoplasm mean fluorescence intensity ratios. Data represent mean ± SEM from three biological replicates (>10,000 PVs each; ∼16 wells/replicate). Statistical significance was determined One-way ANOVA with Tukey’s multiple comparisons test; p-values are shown above the bars. The dashed line at a ratio of 1 indicates little to no enrichment at the PV versus the host cytoplasm.

### TgGRA8 is proximal to ALG-2, ALIX, and TSG101 at the PVM

Next, we reasoned that if TgGRA8 interacts with host ESCRT proteins then it should be proximal to such proteins at the PVM. To investigate the spatial relationship of TgGRA8 with ALG-2, ALIX, and TSG101, we performed super resolution structured illumination microscopy (SIM) on ME49 infected GFP-TSG101 HeLa cells transfected with ALG-2-HA. SIM imaging revealed that TgGRA8 localized near ALG-2 and TSG101 at multiple regions of the PVM **(Fig 5A)**. Notably, ALG-2 and TSG101 frequently colocalized, consistent with their known interaction (12,13), whereas TgGRA8 was typically adjacent to these ESCRT components. In the magnified insets, a few regions within the PVM displayed clear colocalization of all three proteins TgGRA8, ALG-2, and TSG101. To define the spatial relationship between TgGRA8, ALG-2, and ALIX at the PVM we examined WT HeLa cells transfected for mCherry-ALG-2 and infected with Pru::HA-GRA8 parasites. Compared to ALG-2, fewer sites on the PVM showed colocalization between ALIX and TgGRA8 **(Fig 5B)**, but TgGRA8 was often nearby such sites. Notably, all three proteins colocalized at a few distinct regions on the PVM suggesting potential areas of coordinated interaction, although these were less prominent than those for TgGRA8 and ALG-2 as a pair. TgGRA8 also colocalized with ALIX and TSG101 **(S5 Fig)**. Distinct foci of overlap between ALIX and TSG101 were evident at the PVM suggestive of their cooperative recruitment at discrete locations of PVM. Together, these findings support a working model in which TgGRA8 facilitates or stabilizes the association of ESCRT components at the PVM, thereby contributing to host-parasite interface modulation.

**Fig 5.**
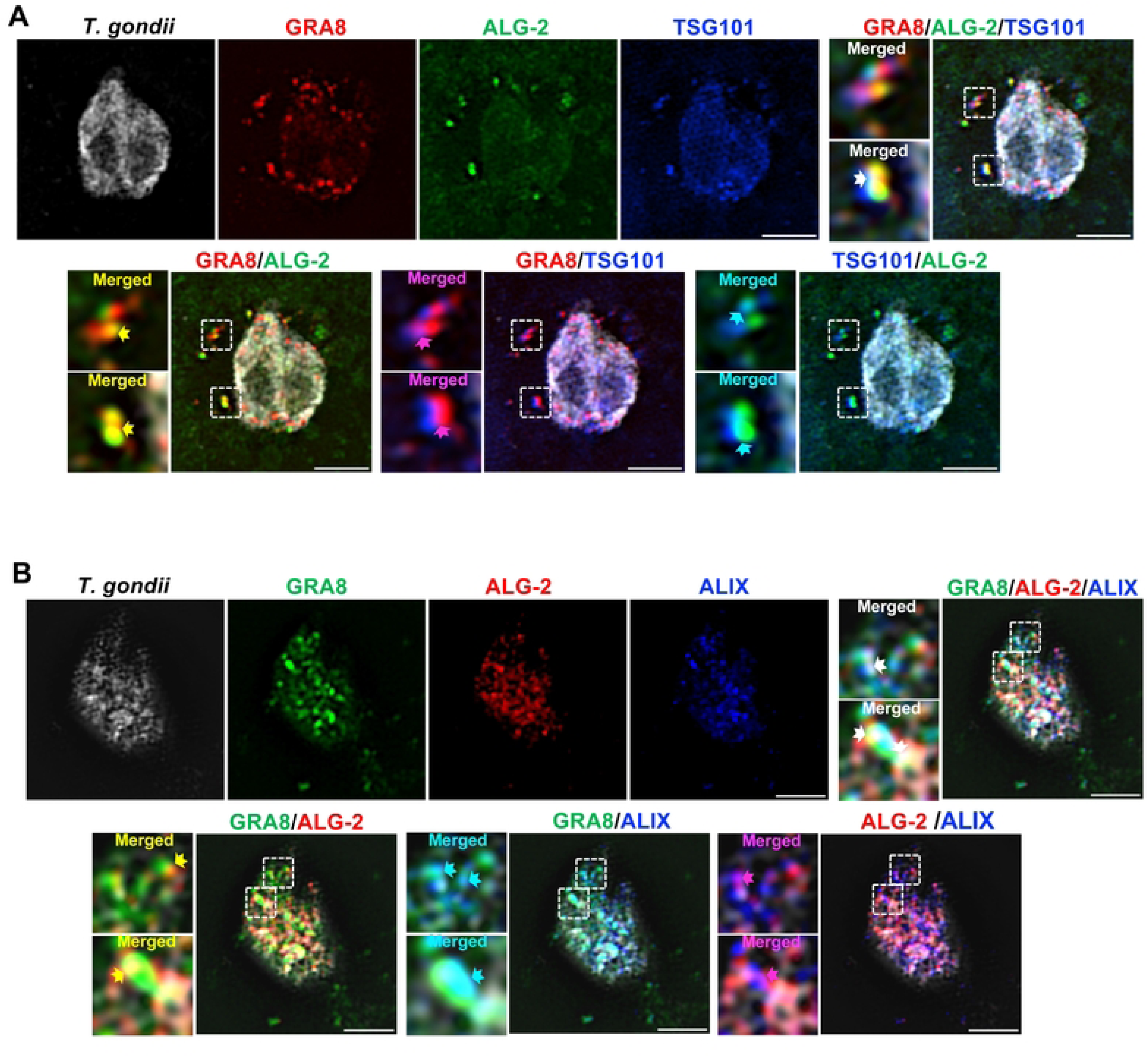
TgGRA8 is proximal to ALG-2, ALIX, and TSG101 at the PVM. **(A, B)** SIM images showing the spatial relationship of TgGRA8 with host ESCRT proteins at the PVM. **(A)** GFP-TSG101 expressing HeLa cells transiently transfected with ALG-2-HA and were infected with ME49 parasites. TgGRA8 was probed using SIM imaging revealed that TgGRA8 (red) localized adjacent to ALG-2 (green) and TSG101 (blue) at multiple regions along the PVM. Merged images and magnified insets of overlap regions (arrows) are shown for TgGRA8-ALG-2, TgGRA8-TSG101, and where all three proteins colocalize. **(B)** HeLa cells expressing mCherry-ALG-2 were infected with Pru::HA-GRA8 parasites and stained for ALIX (green) and TgGRA8 (blue). SIM analysis revealed partial colocalization of TgGRA8 with ALIX, with occasional foci showing triple overlap (arrows). Parasites were visualized using anti-*Toxoplasma* antibodies. Scale bar, 2 µm.

### Identification of conserved motifs in TgGRA8 orthologs

To investigate the conservation of GRA8 across tissue cyst forming coccidian parasites, we used BLAST to identify orthologs of TgGRA8 in *Hammondia hammondi* (*H. hammondi*), *Neospora caninum* (*N. caninum)*, *Besnoitia besnoiti* (*B. besnoiti*), and *Cystoisospora suis* (*C. suis*). Multiple sequence alignment revealed conserved motifs including those that have the potential to bind ALG-2 pocket 1 (PPxY), pocket 2 (MP), and pocket 3 (PPPPGF) **(Fig 6A)**. In the N- and central regions of TgGRA8, these conserved motifs reside within each of 3 tandem repeats, which we termed ALG-2-binding elements (ABE1, ABE2, and ABE3). Comparative analysis showed that NcGRA8 and CsGRA8 contain two additional ABEs inserted between ABE-1 and ABE-2, implying these proteins have evolved toward an increased capacity for binding ALG-2. All GRA8 orthologs harbor a predicted transmembrane domain near their C-terminus, consistent with GRA8 being an integral membrane protein (35). To determine if the TgGRA8 ABE’s are oriented toward the host cytosol, we fixed HFF cells infected with Pru::HA-GRA8 parasites, semi-permeabilized them with saponin, and stained for HA and the parasite surface protein TgSAG1. Some infected cells were positive for both HA and SAG1 indicating permeabilization of both the host plasma membrane and the PVM **(Fig 6B)**. However, other infected cells were only positive for HA, indicating selective permeabilization of the host plasma membrane but not the PVM. Since HA is fused to the ABE-containing N-terminus of TgGRA8, these findings indicate that the ABEs are oriented toward the host cytosol, as shown in **Fig. 6C**. To gain more insights into the molecular basis of the interaction between TgGRA8 and des3-23 ALG-2 (i.e., lacking the unstructured N-terminus), we generated an AlphaFold structural model of a monomeric unit of ALG-2 in complex with the TgGRA8 ABE1 peptide in presence of Ca^2+^ ions. **(Fig 6D)**. AlphaFold multimer modeling of the des3-23 ALG-2-TgGRA8 complex generated a high-confidence model with an interface predicted TM-score (ipTM) of 0.82 and a predicted TM (pTM) of 0.89. The modelling revealed a well-defined binding interface consistent with previously characterized ALG-2 interaction sites with binding protein partners (3). The model predicts that ALG-2 Pockets 1 (blue) and 2 (red) binds to the PPYP (blue) and MP (red) motifs of the TgGRA8 ABE1 peptide. In contrast, ALG-2 Pocket 3 (green) is unoccupied because the intervening sequence prior to the TgGRA8 PPPPGF (green) motif is too short for the motif to reach Pocket 3 of the same ALG-2 molecule. Instead, the TgGRA8 putative pocket 3 binding motif is spatially displaced and potentially available to bind other ALG-2 molecules. Collectively, these findings suggest that TgGRA8 and its orthologs in other tissue cyst-forming coccidians mimic endogenous ALG-2 ligands, enabling their interaction with ALG-2 and potentially modulating ALG-2’s association with downstream ESCRT components.

**Fig 6.**
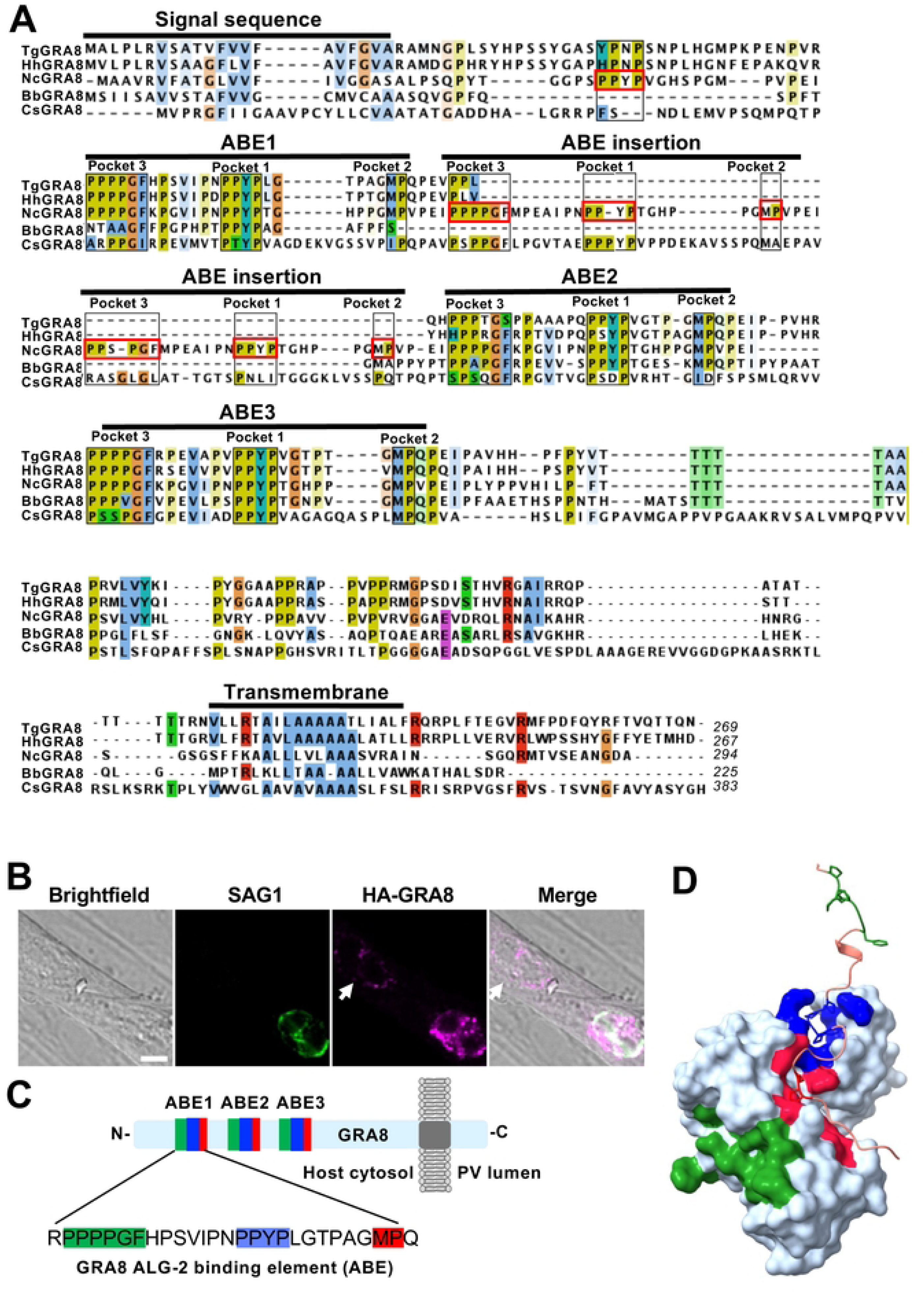
Identification of conserved motifs in TgGRA8 orthologs that are exposed to host cytosol. **(A)** MSA of *T. gondii*, *H. hammondi*, *N. caninum*, *B. besnoiti*, and *C. suis* GRA8 proteins was performed using MAFFT v7 with the L-INS-i algorithm and visualized in Jalview v2.11. Residues exhibiting ≥30% conservation and ≥30% sequence identity are highlighted according to the ClustalX color scheme. The N-terminal signal peptide and C-terminal transmembrane domain are indicated in the alignment. Three ALG-2-binding element (ABE) motifs (ABE1-ABE3) and their associated proline-rich “pocket” motifs (PxxP, PPxY, and PPPPGF) are outlined in the alignment. **(B)** IFA of tachyzoite vacuoles 24 h post-infection showing N-terminally HA-tagged TgGRA8. Semi-permeabilization was performed with 0.01% saponin. The TgGRA8 signal is localized to the PVM, with its N-terminus towards host cytosol defined by negative SAG1 staining. Scale bar: 10µm. **(C)**The key hydrophobic binding pockets of ALG-2 (Pockets 1-3) are highlighted in blue, red, and green, respectively, along with the corresponding interacting residues in the TgGRA8 peptide. **(D)** AlphaFold3 surface model of des3-23 ALG-2 depicting the predicted interaction with the TgGRA8-ABE peptide.

### GRA8 engages ALG-2 through high-affinity interactions

To define the binding kinetics of TgGRA8 and ALG-2, we performed bio-layer interferometry (BLI) experiments with or without Ca^2+^ using synthetic peptides from TgGRA8 ABE1 and recombinant des3-23 ALG-2 (referred as rALG-2^WT^ from here on). Synthetic TgGRA8 peptides corresponding to those for binding to ALG-2 Pocket 1&2 (GRA8-Poc1,2) and Pocket 3 (GRA8-Poc3) were tested alongside an ALG-2 Pocket 1&2-binding peptide from ALIX as a positive control (11). Notably, the TgGRA8 peptides bound to rALG-2 ^WT^ with affinities (K_d_= 2.90 nM and 3.34 nM, respectively) that were two orders of magnitude stronger than the ALIX peptide (K_d_= 660 nM) **(Fig 7A).** A previous study measured a K_d_ of 11 μM for ALIX peptide binding to des3-23 ALG-2 (37), a difference potentially attributable using another approach (plasmon surface resonance). Regardless, as expected, we found that binding of all three peptides to rALG-2^WT^ was Ca²⁺-dependent. Remarkably, a synthetic peptide encompassing the entire TgGRA8 ABE1 bound to rALG-2^WT^ with extremely high affinity (K^d^ < 0.001 nM) that unexpectedly lacked Ca²⁺-dependence. To determine the pocket specificity, we tested binding of TgGRA8 peptides to recombinantly expressed des3-23 ALG-2^ΔGF^ (referred as rALG-2^ΔGF^ from here on), an alternatively spliced variant that lacks two key residues in pocket 1&2 but retains a functional pocket 3. As expected, the GRA8-Poc1,2 peptide and the ALIX peptide showed no binding to rALG-2^ΔGF^, whereas GRA8-Poc3 bound to rALG-2^ΔGF^ in a Ca²⁺-dependent manner **(S6 Fig)**. Interestingly, GRA8-FL bound to rALG-2^ΔGF^ with high affinity (K_d_ = 0.19 nM) and was partly Ca²⁺-dependent (K_d_= 3.33 nM). Together these findings confirm that TgGRA8 engages all three binding pockets of ALG-2 and that these interactions have unique properties compared to other ALG-2 binding partners.

**Fig 7.**
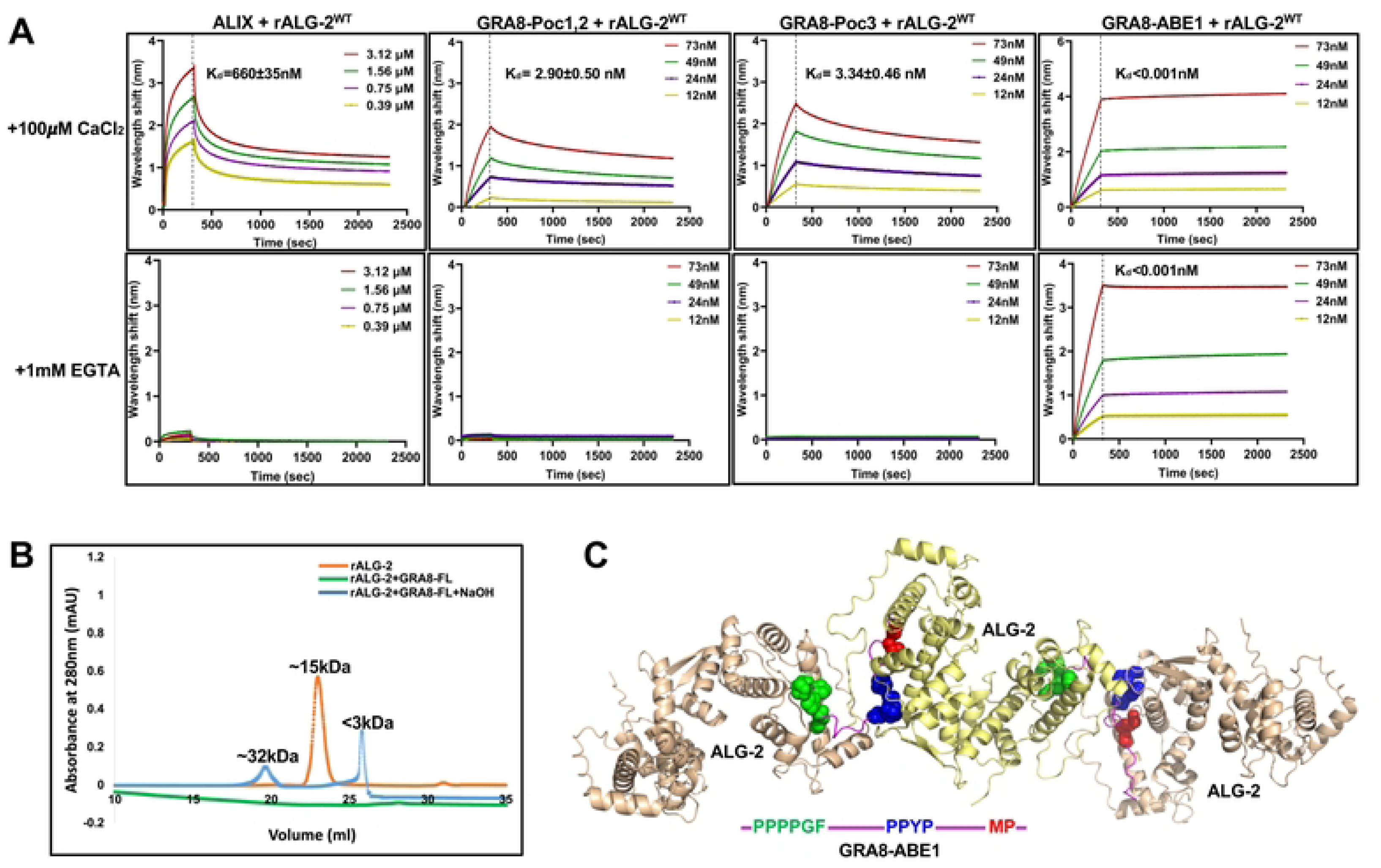
High-affinity binding of the TgGRA8-ABE1 peptide to rALG-2^WT^ promotes the formation of higher-order assemblies. **(A)** BLI assay showing binding affinity of N-terminally biotinylated ALIX, GRA8-Poc1,2, GRA8-Poc3, and GRA8-FL peptides, which was immobilized onto streptavidin biosensors. The analysis involved exposing these peptides to an array of different concentrations of des3-23ALG-2^WT^ (concentrations shown in the key) in a solution containing 100 µM Ca^2+^ or no added Ca^2+^ and 1 mM EGTA. The association between the peptides and proteins for both the experiments were measured for 300 s followed by dissociation for 2,000s. Data shown here are from 2 independent experiments conducted for each condition. BLI sensorgram were baseline-corrected and globally fitted using a 1:1 binding model. **(B)** Affinity-purified rALG-2^WT^ protein alone or mixed with synthetic TgGRA8-ABE1 at a 1:1 equimolar ratio, was analyzed by size-exclusion chromatography under reducing conditions (+β-ME) to assess peptide-protein complex formation. The column was subsequently flushed with 0.5N NaOH to remove any bound complex. **(C)** AlphaFold modeling of two TgGRA8 ABE1 peptides with three ALG-2 dimers predicts that TgGRA8 ABE1 bridges adjacent ALG-2 molecules, forming multimeric assemblies.

To further investigate the unique properties of GRA8-FL binding to rALG-2^WT^, we performed size-exclusion chromatography (SEC) experiments. rALG-2^WT^ alone eluted as a single ∼15 kDa peak **(Fig 7B).** However, rALG-2^WT^ mixed with GRA8-FL failed to elute from the column, implying formation of a large complex. Accordingly, treating the column with 0.5N NaOH resulted in elution of two peaks sized ∼32kDa and <3kDa. We reasoned based on our earlier structural modeling **(Fig 6)** that the spatially displaced GRA8-FL pocket 3 binding motif could be available to bind other ALG-2 molecules to form a super complex. To test this, we used Alphafold multimer to model two molecules of TgGRA8-ABE1 with 6 molecules of ALG-2 (i.e., 3 homodimers). This analysis predicted the formation of a supramolecular assembly (ipTM: 0.49 and pTM: 0.55) driven by TgGRA8’s three distinct ABMs (Poc1: blue, Poc2: red, Poc3: green), enabling it to coordinate multiple ALG-2 dimers through its multivalent topology **(Fig 7C)**. Within the model, the TgGRA8 Pocket 1 and Pocket 2 motifs interact with the corresponding binding sites on one ALG-2 monomer, while the partner ALG-2 monomer remains unoccupied and thus available for association with host ESCRT components. The spatially displaced TgGRA8 pocket 3 motif is predicted to occupy pocket 3 of an adjacent ALG-2 molecule, producing an extended multimeric bridging architecture. Development of supramolecular complexes on the BLI probe could have led to inaccurate measurements, potentially explaining the unexpected Ca²⁺-independent interaction between TgGRA8-ABE1 and rALG-2^WT^.

### Multiple pockets of ALG-2 mediate Ca^2+^-dependent recruitment to the *Toxoplasma* PV

To identify functional determinants of ALG-2 required for recruitment to the *Toxoplasma* PV, we measured recruitment of mCherry-ALG-2^WT^ versus that of the ALG-2^F85A^ (Poc3 binding mutant) (3), ALG-2^ΔGF^ (Pocket 1 binding mutant) (7,8), ALG-2^ΔGF/F85A^ (Poc1&3 binding mutant), and the ALG-2^E47A/E114A^ (Ca²⁺-binding mutant) in transiently transfected HeLa cells infected with type I (RH) or type II (ME49) parasites. In HeLa cells infected with the type I RH strain, individual mutation of either pocket 1 or pocket 3 had only modest effects on ALG-2 recruitment to the PV. In contrast, combined mutation of both pockets 1 and 3, or mutation of the Ca²⁺-binding residues, markedly reduced ALG-2 abundance at the PV. **(Fig 8A, B)**. In contrast, infection with the type II ME49 strain resulted in a more pronounced reduction in PV recruitment of all the ALG-2 mutants **(Fig 8C, D).** Analysis of well-level data confirmed the differences observed for the biological replicates (**S8 Fig A, B)**. Overall, these results suggest that ALG-2 recruitment to the PV requires Ca^2+^-dependent interactions with multiple pockets of ALG-2 in a manner that is influenced by the strain of the parasite.

**Fig 8.**
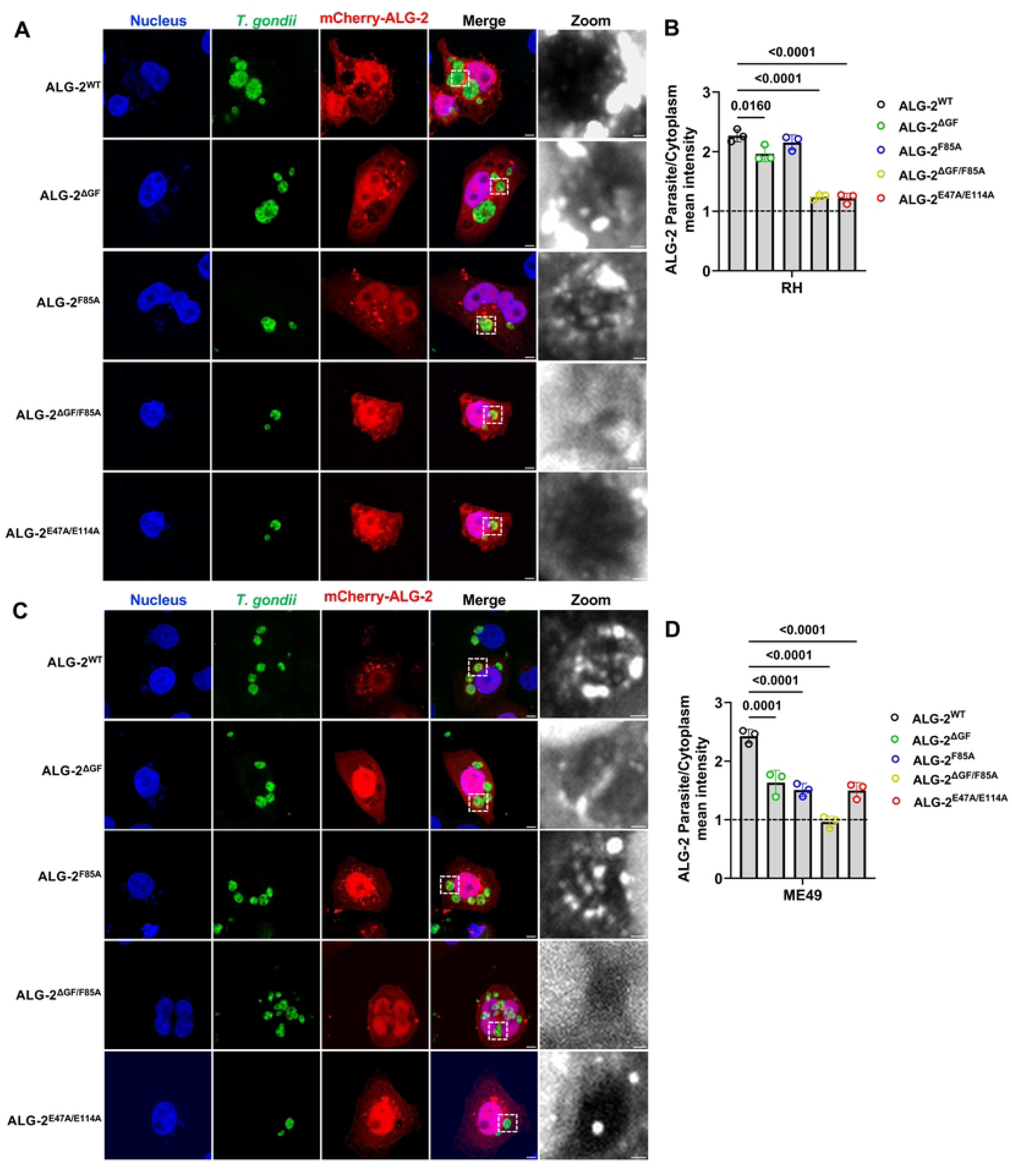
TgGRA8 binds directly to ALG-2 with high affinity in a Ca^2+^-dependent manner. **(A, B)** ALG-2 and TgGRA8 interact in pocket specific manner. HeLa cells were transfected with mCherry-ALG-2^WT^ or mCherry-ALG-2^F85A^ or mCherry-ALG-2^ΔGF^ or mCherry-ALG-2^ΔGF/F85A^ or mCherry-ALG2^E47A/E114A^ and infected with either RHΔku80WT **(A, B)** or ME49Δku80WT **(C, D)** parasites to study the recruitment of ALG-2. After 24 h of infection cells were fixed and stained to assess ALG-2 recruitment to the PV. Representative IFA images **(A, C)** and quantification **(B, D)** show the recruitment of WT and mutant ALG-2 to the PVM, expressed as the mean fluorescence intensity ratio (PVM-to-cytosol) across 3 independent experiments (>10,000 PVs per replicate). Scale bar, 10 µm. Black and white zoomed insets representing the ALG-2 recruitment on the PVM. Scale bar, 2 µm. Each data point represents the well average (∼16 wells per biological replicates) ± SEM. Statistical analysis was performed using one-way ANOVA with Tukey’s multiple comparisons test. The dashed line at a ratio of 1 indicates little to no enrichment at the PV versus the host cytoplasm.

### Parasites lacking GRA8 have an abnormal metabolic profile like that of ingestion mutants

To delineate the functional metabolic consequences of GRA8 ablation, we performed targeted metabolomics on RHΔ*ku80*WT, RHΔ*ku80*Δ*gra8*, and RHΔ*ku80*Δ*gra8GRA8* (GRA8 complement) and compared the findings to datasets from two ingestion mutant strains, RHΔ*gra14* and RHΔ*cpl* (32). Principal component analysis (PCA) of normalized metabolite abundances revealed clear strain-dependent clustering, with Δ*gra8* clustering tightly with Δ*gra14*, which together overlapped with a more dispersed Δ*cpl* cluster (**Fig 9A**). Data from GRA8 complemented parasites clustered with that of the parental strains, indicating that restoration of GRA8 expression resulted in a normal metabolic profile. Quantified metabolites were classified into six major biochemical categories: amino acids, nucleosides, glycolytic intermediates, tricarboxylic acid (TCA) cycle metabolites, pentose phosphate pathway (PPP) components, and energy-carrying molecules (ATP, ADP, NAD⁺, NADP⁺). Intriguingly, the metabolic profile of Δ*gra8* parasites (**S3 Data**) exhibited striking similarity to that of the Δ*gra14* ingestion mutant (**Fig 9B; S7A Fig**), suggesting that these two dense granule effectors converge on shared metabolic regulatory nodes or participate in overlapping nutrient acquisition pathways. Consistent with these metabolite-level changes, pathway enrichment analysis identified purine metabolism, amino acid metabolism, and central carbon metabolism as major pathways altered upon loss of GRA8, GRA14, or CPL **(Fig 9C; S7 B,C Fig).** Together, these analyses indicate that GRA8 ablation induces broad metabolic remodeling in *T. gondii*, affecting mainly nucleotide, amino acid, and central carbon metabolic pathways. Also, whereas assessing ingestion directly is difficult because of inter-experiment variability, metabolomics analysis is more robust and informative of global effects on parasite physiology. Notably, parasites lacking GRA8 showed no generalized fitness defect in the Giuliano et al. genome-wide mouse CRISPR screen and retained positive fitness in cell culture (mean fitness score, +1.82) (38). Thus, the metabolic defects caused by GRA8 ablation likely reflect a specific need for GRA8 in parasite metabolic homeostasis rather than impaired baseline replication.

**Fig 9.**
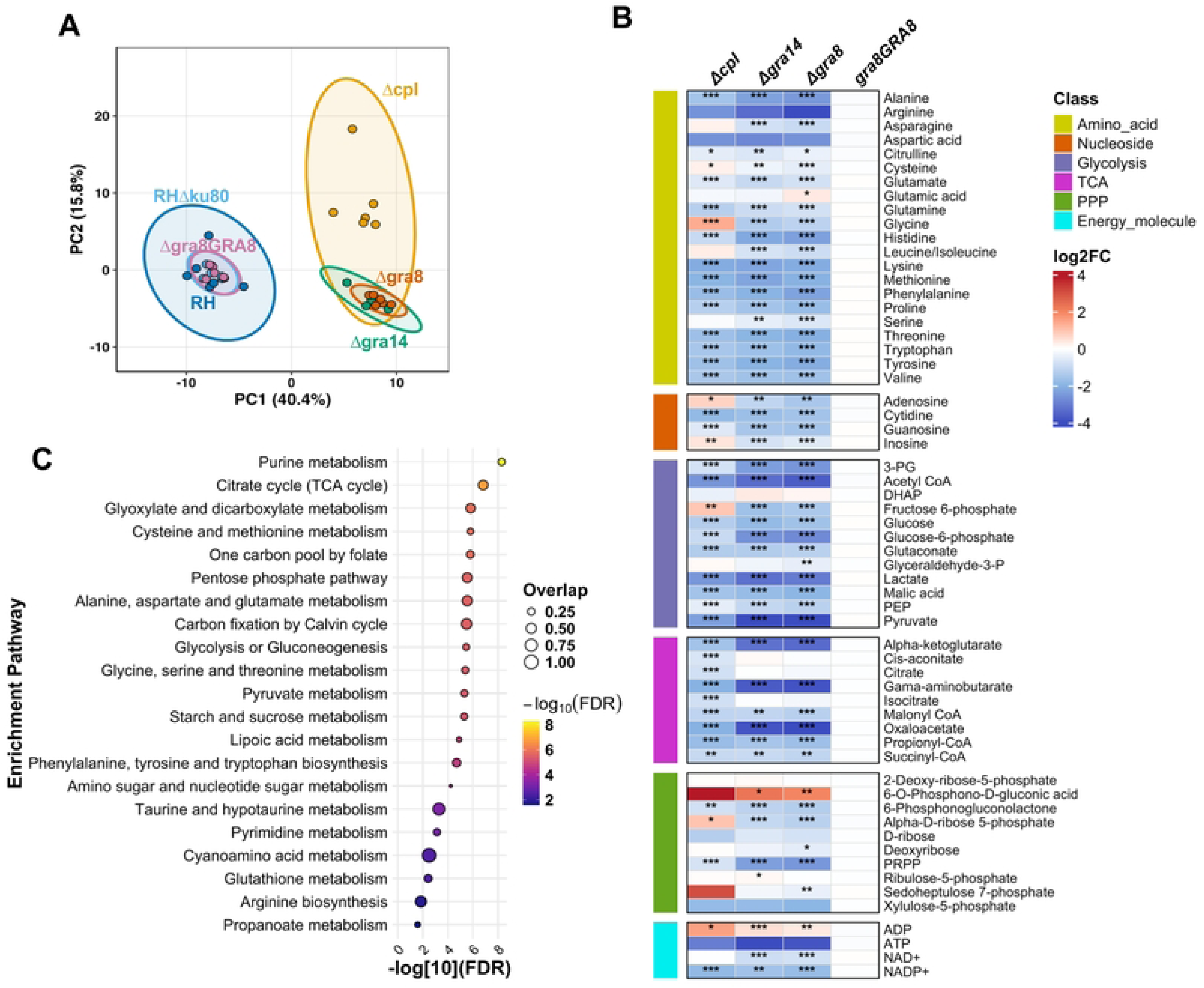
TgGRA8 ablation drives broad metabolic reprogramming converging on glycolytic, TCA cycle, and bioenergetic pathway suppression. **(A)** Principal component analysis (PCA) of normalized metabolite abundances from RHΔ*ku80*WT, RHΔ*ku80*Δgra8, RHΔ*ku80*Δ*gra8*::*GRA8* (Δ*gra8GRA8*), RHΔ*gra14*, and RHΔ*cpl* parasite strains. Each data point represents an independent biological replicate. Shaded ellipses denote 95% confidence intervals for each strain grouping. **(B)** Heatmap showing log2 fold changes in metabolite abundance relative to WT parasites for Δ*cpl*, Δ*gra14*, and Δ*gra8* strains. Metabolites are grouped by biochemical class, denoted by colored sidebar annotations: amino acids (yellow green), nucleosides (orange red), glycolysis intermediates (light purple), TCA cycle metabolites (magenta), pentose phosphate pathway (PPP), intermediates (olive green), and energy molecules (cyan). Color scale ranges from blue to red (log2FC = −4 to +4). Asterisks indicate statistical significance of pairwise comparisons relative to WT: *= 0.05, **= 0.01, ***=0.001. t-test. n = 6 biological replicates per strain. **(C)** Pathway enrichment analysis was performed using MetaboAnalyst (69) on metabolites altered in RHΔku80Δgra8 parasites relative to RHΔku80WT. Enriched pathways are plotted by enrichment factor. Point color indicates significance as -log10(FDR), and point size indicates pathway impact.

## Discussion

In this study, we uncover a novel mechanism by which *T. gondii* dense granule effector TgGRA8 co-opts the host ESCRT adaptor ALG-2 at the PVM during intracellular replication. ALG-2 pulldown assays revealed TgGRA8 as a significantly enriched effector, while reciprocal TgGRA8 co-IPs identified multiple host ESCRT-associated proteins. Functional and imaging analyses demonstrated that TgGRA8 is essential for ALG-2 and ALIX recruitment to the PVM in both type I (RH) and type II (ME49) parasites. Parasites lacking TgGRA8 exhibited marked metabolic perturbations, suggesting a role in host nutrient acquisition.

Host ESCRT machinery is critical for ‘reverse-topology’ membrane scission that severs the necks of the membranes that bud away from the host cytosol (39). The process of sequential recruitment of ESCRT-I/ALIX, ESCRT-III polymerization, and VPS4-driven disassembly, drives multivesicular endosome formation and supports cytokinesis, viral budding, and plasma membrane repair (40). Enveloped viruses such as HIV-1 and Ebola virus hijack this machinery to complete virion budding, primarily via PTAP (TSG101-binding) and YPXL (ALIX-binding) late-domain motifs in their matrix proteins including HIV-1 Gag (20,41–43). In an example of convergent evolution, *Toxoplasma* uses a PTAP motif in TgGRA14 to recruit TSG101 to its PVM for ingestion of host cytosolic material and, together with TgGRA64, for PV entrapment of lipid rich Rab11a positive vesicles derived from the endocytic recycling compartment (29,30). Prior work showed that the TgGRA14 C-terminal domain can functionally substitute for the late domain of HIV-1 Gag to produce virus like particles (VLPs) in an ESCRT-dependent manner and that the TgGRA14 PTAP and YPXL motifs were necessary for VLP production (29). However, this same study found that whereas the TgGRA14 PTAP motif is necessary for TSG101 recruitment to the PV, neither the expression of TgGRA14 nor mutation of the YPXL motif affected PV recruitment of ALIX. Herein we detected a modest but significant reduction in ALIX recruitment upon deletion of TgGRA14, a finding that is likely attributed to the more robust quantification afforded by using high content imaging and semi-automated image analysis. This result also aligns with the TgGRA14 YPXL motif contributing to VLP production in the functional substitution experiments (29). We likewise detected a modest contribution of TgGRA64 to the PV recruitment of ALIX. Although TgGRA64 possesses a poly-arginine motif analogous to the ALIX-interacting region of HSV-1 UL31 that drives ESCRT-III recruitment (34,44), whether this motif is functional for binding ALIX remains to be determined. Regardless, whereas our findings suggest both unique and additive roles for TgGRA14 and TgGRA64 in the PV recruitment of TSG101 and ALIX, these proteins do not account for the PV recruitment of ALG-2.

ALG-2 immunoprecipitation and gene knockout experiments identified TgGRA8 as a critical determinant of ALG-2 recruitment to the PV. TgGRA8 encodes a 267-amino acid protein with an N-terminal signal peptide, a central region containing degenerate proline-rich repeats, and a C-terminal transmembrane domain (35). A previous study suggested that TgGRA8 is imported into host mitochondria where it interacts with components of the ATP synthase F(1) complex and the mitochondrial matrix protein Sirtuin-3 (45). However, this prior work studied TgGRA8 alone in host cells, either by adding the recombinant protein to cells exogenously or by expressing it via transfection. Herein, we failed to identify members of the ATP synthase F(1) complex or Sirtuin-3 in our immunoprecipitation experiments using lysates from infected cells. Instead, we found that TgGRA8’s interaction network includes direct binding to ALG-2 along with enrichment of other host ESCRT proteins. Sequence analysis revealed that TgGRA8 harbors three ALG-2-binding elements (ABE1-3) within its N-terminal region exposed to the host cytosol. Remarkably, these ABEs encompass ABM-1 and ABM-2, an arrangement not previously seen in any ALG-binding proteins. BLI experiments using TgGRA8-derived peptides corresponding to ABM-1 (ALG-2 pocket 1 and 2) or ABM-2 (ALG-2 pocket 3) demonstrated Ca²⁺-dependent binding to ALG-2, confirming that TgGRA8 can engage ALG-2 through multiple binding modes. GRA8 ABM-2 was also was able to interact with ALG-2^ΔGF122^ splice variant previously reported to be unable to interact with ALIX (7,8,46) indicating that TgGRA8 can engage ALG-2 through alternative binding pockets. Unexpectedly, GRA8-FL bound ALG-2 in a Ca^2+^-independent manner. The GRA8-FL-ALG-2 interaction exhibited rapid association and essentially no measurable dissociation over extended wash phases, indicating formation of an unusually stable complex. The apparent Ca^2+^-independence and failure to dissociate might be manifestations of the multivalent interactions predicted by AlphaFold modeling. Structural predictions suggest that TgGRA8 might bridge ALG-2 dimers via spatially distinct ABEs, forming higher-order assemblies that could stabilize ALG-2 at the PVM and strengthen ESCRT localization during infection. Nevertheless, abolishing ALG-2’s ability to bind Ca^2+^ eliminated its association with the PV, indicating that ALG-2 recruitment to the PV is Ca^2+^-dependent. Precisely how ALG-2 is activated by Ca^2+^ at the PVM and the source of Ca^2+^ for such activation remain to be determined.

The ability of ALG-2 to form homodimers bestows upon it the ability to simultaneously engage multiple ESCRT proteins such as TSG101 and ALIX that contain ABM-1 motifs for pocket 1 and pocket 2 (9–11,47,48). Our data demonstrates that TgGRA8 engages ALG-2 through multiple, topologically distinct binding to all three binding pockets of ALG-2. Structural modeling indicates that the dimeric structure of ALG-2 allows for ALG-2 binding to other proteins even when it is engaged with TgGRA8, which is consistent with TgGRA8 not only co-immunoprecipitating with ALG-2 but also with many other ESCRT proteins including TSG101 and ALIX. As Sec31A was not detected in the pulldown experiments this might reflect a measure of specificity in ALG-2’s partnerships while engaged with TgGRA8. Collectively, our findings suggest that TgGRA8 primarily promotes ALG-2 and ALIX recruitment, while TgGRA14 serves as the dominant recruiter of TSG101. We therefore propose a model in which TgGRA8 acts as a multivalent adaptor that stabilizes ALIX-TSG101 assemblies by positioning Ca^2+^-bound ALG-2 dimers at discrete sites on the PVM. Super-resolution imaging supports this concept, revealing a striking proximity of ALIX, ALG-2, and TSG101 at the PVM, either overlapping with TgGRA8 or forming adjacent puncta. Functional assays indicate that disruption of individual ALG-2-binding pockets only modestly affects ALG-2 recruitment in type I parasites, suggesting that TgGRA8 can compensate for the loss by alternative ALG-2-binding motifs. By contrast, type II parasites show stronger dependency on specific pocket interactions, implying strain-specific preferences for ALG-2 engagement. Whether these strain dependencies are driven by GRA8 directly or by differences in how GRA8 is displayed at the PVM remains to be determined. Regardless, ALG-2 binding to Ca^2+^ was necessary for recruitment in both strains, consistent with the requirement of Ca^2+^ for stabilizing ALG-2 dimers and enabling their engagement with target motifs. This suggests that TgGRA8 may reinforce ALG-2 localization at the PVM only when the ALG-2 dimer is in its Ca^2+^-activated state, whereas the absence of Ca^2+^ likely limits TgGRA8’s ability to engage multiple binding surfaces simultaneously.

This Ca^2+^ dependence is consistent with established roles for ALG-2 as a prototypic PEF Ca^2+^ sensor that functions as a damage-responsive adaptor at membranes (4,11,49). During plasma membrane injury, extracellular Ca^2+^ influx triggers ALG-2 activation, driving recruitment of ALIX, ESCRT-III components, and VPS4 to the wound site, mediating membrane scission and shedding of damaged membrane domains to restore plasma membrane integrity (50). A parallel Ca^2+^-dependent ALG-2-ESCRT axis has been delineated at lysosomes, where membrane damage or osmotic stress induces local Ca^2+^ release, triggering recruitment of ALG-2 and its associated ESCRT machinery. Previous *in vitro* membrane reconstitution studies have shown that Ca^2+^ - bound ALG-2 is sufficient to target ALIX to negatively charged membranes and promote sequential assembly of ESCRT-III subunits together with the AAA⁺ ATPase VPS4B, establishing ALG-2 as an upstream Ca^2+^-responsive organizer of lysosomal membrane repair (51). More recent work extends this paradigm by demonstrating that lysosomal Ca^2+^ release can precondition lysosomes through ALG-2-dependent ESCRT recruitment before overt membrane rupture, indicating that Ca^2+^-activated ALG-2-ESCRT assembly functions not only as a repair response to membrane damage, but also as a preventative mechanism that enhances lysosomal membrane resilience against osmotic stress (52). This Ca^2+^-dependent function of ALG-2 in organizing ESCRT machinery at stressed intracellular membranes provides a mechanistic basis for interpreting its recruitment to the *T. gondii* PVM as a potential site of localized Ca^2+^-activated ALG-2-ESCRT engagement. Through hydrophobic pocket-mediated recognition of proline-rich motifs, dimeric ALG-2 can stabilize or bridge membrane-associated ESCRT complexes, including ALIX and TSG101 (4,11,13,53). Thus, by recruiting ALG-2 to the PVM, GRA8 may create a membrane-proximal ALG-2 docking platform that promotes localized ALIX/ESCRT recruitment. By analogy with injured plasma membranes and stressed lysosomes, the *T. gondii* PVM may represent a specialized host-pathogen membrane interface where local Ca^2+^ responsive ALG-2 scaffolding organizes ESCRT machinery, potentially contributing to vacuolar membrane remodeling, PVM maintenance, and parasite nutrient acquisition. Precisely how ALG-2 is activated by Ca^2+^ at the PVM and the source of such Ca^2+^ should be the subject of future investigation.

*T. gondii* displays marked metabolic flexibility that likely underlies its ability to proliferate across diverse host cell types and nutrient environments (reviewed in (54). Across life stages, tachyzoites balance *de novo* biosynthesis with extensive salvage and import, and remodel central carbon metabolism (glycolysis, the TCA cycle, and the pentose phosphate pathway) to maintain redox balance and biomass precursor supply when key inputs are limiting. Prior work identified metabolic signatures of disrupted parasite ingestion, including mutants lacking TgGRA14, TgCPL (32), or a functional micropore (55), the likely site of ingestion. Here, we find that TgGRA8 ablation produces metabolic changes that mirror these ingestion-pathway mutants. Although ingestion is difficult to quantify, metabolomics provides a robust readout consistent with altered access to host resources. Coordinated disruption of amino acid, purine, and central carbon pools without overt growth defects may reflect nutrient-replete culture conditions and/or compensatory reliance on alternate acquisition routes (e.g., via PVM pores) and parasite-intrinsic biosynthesis. Despite this adaptability, *Toxoplasma* is auxotrophic for several metabolites and depends on host uptake and salvage. Amino-acid acquisition is critical, with multiple essential amino acids requiring host-derived salvage. *Toxoplasma* also cannot synthesize purines *de novo* and relies on purine salvage for nucleotide production (56,57); thus, reduced access to host nucleosides/nucleobases can deplete purine-associated metabolites and drive compensatory shifts such as increased PPP activity to support ribose-phosphate production under precursor limitation, as seen in the TgGRA14 and TgCPL ingestion mutants and TgGRA8-deficient parasites. Together, these auxotrophies highlight that parasite survival depends on robust nutrient import/salvage and host-interface processes that maintain access to amino acids and nucleosides needed for growth and replication.

## Conclusions

Together, our study identifies TgGRA8 as a dense granule effector that promotes ALG-2-dependent ESCRT recruitment to the *T. gondii* PVM by engaging through ALG-2-binding elements (ABE1-3). TgGRA8 functions alongside other dense granule effectors, particularly TgGRA14 and TgGRA64, to coordinate distinct arms of host ESCRT recruitment, with TgGRA8 primarily supporting ALG-2/ALIX engagement and TgGRA14 serving as a dominant recruiter of TSG101. Our biochemical and imaging data supports a non-canonical, Ca^2+^-dependent mode of ALG-2 engagement where TgGRA8 potentially bridges multiple copies of ALG-2 at the PV. The conservation of GRA8 ABEs among tissue cyst–forming coccidians suggests that this mechanism may represent a lineage-specific adaptation associated with parasite persistence. Metabolomic profiling further shows that loss of GRA8 produces broad defects in amino acid, purine, and central carbon metabolism that overlap with the RHΔ*gra14* ingestion mutant, implicating TgGRA8 in parasite metabolic homeostasis and host-derived nutrient acquisition. Thus, TgGRA8 links ESCRT organization at the PVM to parasite metabolic homeostasis, suggesting a role in coordinating host membrane remodeling with nutrient acquisition during intracellular infection.

## Materials and Methods

### Host cell culture and transfection

All host cell lines, including human foreskin fibroblast (HFF), HeLa, and GFP-TSG101 HeLa cells, were grown in Dulbecco’s Modified Eagle’s Medium (DMEM) supplemented with 10% Cosmic Calf Serum, 20 mM HEPES, 5 μg/mL penicillin/streptomycin, and 2 mM L-glutamine (named as D10). HeLa cells were transfected using Lipofectamine 3000 (Thermo Fisher Scientific, Cat # L3000008) following the manufacturer’s instructions. Briefly, cells were seeded in Poly-D-Lysine (Thermo Fisher Scientific, Cat # A3890401) coated 6-well plates at a density of 3.0 × 10^5^ cells per well and transfected with 4 μg of plasmid DNA per well. Transfected cells were incubated for 20-24 h before being subjected to infection with specific *T. gondii* strains.

### Parasite culture and parental strains

*T. gondii* strains were maintained by serial passage in human foreskin fibroblast (HFF). RHΔ*ku80*Δ*hxgprt* and ME49Δ*ku80Δ*hxgprt were used as parental strains for facile genetic manipulation (58,59).

### Generation of the RHΔ*gra14,* RHΔ*gra64* and RHΔ*gra14*Δ*gra64* strains

The RHΔ*gra14* strain is described in (29). The RHΔ*gra64* strain is described in (34). The RHΔ*gra14*Δ*gra64* strain is described in Romano et al. (30). For a full list of primers used for cloning and genetic manipulation in this paper, please refer to **S1 Table**. For all Cas9-mediated genetic manipulations, single guide RNAs (sgRNA) were designed using CHOPCHOP (https://chopchop.cbu.uib.no/) (60) and cloned into the p-HXGPRT-Cas9-GFP plasmid backbone using KLD (Kinase, Ligase, and Dpn1) reactions (New England Biolabs, Ipswich, MA), as previously described (61). Next, 100-bp donor oligonucleotides were designed and synthesized (Thermo Fisher) with homologous arms targeting the region of interest and encoding a stop codon (inserted in place of the start codon for a knockout), HA-tag, or a start codon (inserted in place of the ectopic stop codon in the knockout strain for complement transfection).

### Generation of the RHΔ*gra8* and ME49Δ*gra8* strains

First, CRISPR-Cas9 gene editing was used to target the endogenous TgGRA8 locus by transfecting vector expressing an HXGPRT resistance cassette and Cas9 with the guide RNA sequence *5’-*GTTTCCGCGCTGTAGTAAGA-3’, along with both the forward and reverse 100bp donor oligo, designed to insert a STOP codon (indicated in lower cases) at the N-terminus of the gene for the purpose of generating a gene knockout. Following transfection, 1×10^6^ to 5×10^6^ parasites were electroporated in CytoMix (62) after harvesting egressed parasites from HFF monolayers and filtering through 5-μm filters. Parasites were selected for mycophenolic acid (25 µg/ml) and xanthine (50 µg/ml) for 24 h post transfection for 6 days. Transfected parasites were subcloned by limiting dilution in 96-well plates after sufficient parasite egress had been observed. Δ*gra8* subclones were selected based on negative TgGRA8 antibody (mouse anti-TgGRA8 1:1500, a kind gift from Gary Ward, University of Vermont) (35) and confirmed by sequencing the PCR products amplified from genomic DNA by Sanger sequencing, using the following primers to amplify the 5’UTR and N-terminus region of the *TgGRA8* locus to confirm the insertion of stop codons.

### Generation of RHΔgra8GRA8 and ME49Δgra8GRA8 complement strains

First, CRISPR-Cas9 gene editing was used to target the stop codon in the RHΔ*gra8* and ME49Δ*gra8* strain by transfecting vector expressing an HXGPRT resistance cassette and Cas9 with the guide RNA sequence *5’-TAGCTAACTAGCTAACTGAC-3’*, along with both forward and reverse primer, designed to repair the gene. Transfection, selection, and subcloning was performed as described above. Subclones were selected based on positive TgGRA8 antibody labeling and confirmed by Sanger sequencing using the primers described above.

### Generation of Pru::HA-GRA8

First, CRISPR-Cas9 gene editing was designed to insert a hemagglutinin (HA) sequence at the N-terminus of the gene after the signal peptide to tag the gene for further study by transfecting vector expressing an HXGPRT resistance cassette and Cas9 with the guide RNA sequence 5’-*TGAACGGTCCTTTGAGTTAT*-3’, along with both forward and reverse primer. Transfection, selection, and subcloning was performed as described above. Subclones were selected based on positive HA antibody (Rat anti-HA 3F10, 1:500, Sigma-Aldrich) and confirmed by sequencing the PCR products amplified from genomic DNA by Sanger sequencing, using the following primers to amplify the 5’UTR and N-terminus region of the *TgGRA8* locus to confirm the insertion of the HA tag.

### Immunofluorescence and immunoblotting

HeLa and GFP-TSG101 HeLa cells were washed and fixed with 4% paraformaldehyde (PFA) in 1X PBS for 10 min at RT, permeabilized with 0.3% Triton X-100 for 15 min at RT and further blocked with 10% FBS for 1hr at RT. Primary and secondary antibody staining were performed at 4C and RT for overnight and 1 h in 5% FBS. Imaging was performed in the Nikon N-SIM super-resolution or a Yokogawa CellVoyager 8000 spinning disc confocal high content imaging instrument. Nitrocellulose membranes were blocked in 5% milk, probed with primary antibodies, and then incubated with HRP-conjugated secondary antibodies. For detection, membranes were incubated with either SuperSignal West Pico PLUS Chemiluminescent Substrate (Thermo Fisher Scientific, Cat# 34580) or SuperSignal West Femto Maximum Sensitivity Substrate (Thermo Fisher Scientific, Cat# 34095). Chemiluminescence signal was captured using the Azure 300 Chemiluminescent Imaging System.

### ESCRT protein (ALG-2, ALIX, and TSG101) recruitment

For ALG-2 recruitment experiments, HeLa cells were grown in 6-well well plate and transfected with either pcDNA-ALG-2-HA or mCherry ALG-2 WT or mutant plasmids. The cells were further trypsinized and washed once with D10 and reseeded in a Poly-D-Lysine treated 384 well plate (Revvity Cat# 50-209-9109). The cells were allowed to adhere for 4-6 h and infected with type I strains at an MOI of 4 or type II strains with an MOI of 10 for 24 h. Cells were fixed and stained for ALG-2 using anti-HA antibody (Roche, Cat# 11867423001). For ALIX recruitment, similarly HeLa cells were seeded in 384 well plates and ALIX were stained using purified monoclonal anti-ALIX Ab (BioLegend, Cat# 634502). For TSG101, HeLa cells expressing endogenously tagged GFP–TSG101 were similarly seeded in 384 well plate and in all above parasites were labeled with biotin-conjugated polyclonal anti-*Toxoplasma* antibody (Thermo Fisher Scientific, Cat# PA1-73002), nuclei were stained with Hoechst 33342 (Sigma, Cat# TA9H97BAECD2), and host cell membranes were visualized using HCS CellMask Deep Red (CMDR) stain (Thermo Fisher Scientific, Cat# H32721). Plates were imaged at 40× magnification using a Yokogawa CellVoyager 8000 spinning-disk confocal high-content imaging system. Nine fields of view (FOVs) were captured per well, with Z-stacks spanning 10 µm (1 µm steps) per channel, followed by maximum intensity projection (MIP) generation. Quantitative image analysis was performed by classifying cells (CMDR MIP), nuclei (Hoechst MIP), cytoplasm (cell area excluding nucleus and PV), and parasites (Anti-Toxo MIP). Fluorescence intensities of ALG-2, ALIX or TSG101 were measured at the PVM and in the cytoplasm, and PV-to-cytoplasm intensity ratios were calculated. We excluded unstained and untransduced cells using a threshold defined as the 95th percentile of channel of interest intensity. All quantitative image analysis was performed as described in (33) in detail. For ALG-2 IFA, HeLa cells were cultured on 1.5 mm coverslips in 12-well plates and infected with type I parasites at an MOI of 4 or type II parasites at an MOI of 10 for 24 h. Cells were fixed and permeabilized using the Becton Dickinson Cytofix/Cytoperm Fixation/Permeabilization Kit (Cat# 554714) according to the manufacturer’s protocol. Coverslips were stained with polyclonal anti-*Toxoplasma* antibody (Thermo Fisher Scientific, Cat# PA1-73002), nuclei were stained with Hoechst 33342 (Sigma, Cat# TA9H97BAECD2), and ALG-2 was stained anti-ALG-2/PDCD6 polyclonal antibody (Cat# 12303-1-AP). Coverslips were imaged on a Nikon X1 Yokogawa spinning-disk microscope using a 100X magnification.

### TSG101 UEV domain protein expression and purification

A DNA construct encoding the TSG101 UEV domain (aa 2-145) was cloned into the pGST2 vector (63). This vector results in an N-terminally GST-TEV tagged TSG101 UEV domain. The resulting plasmid was transformed into *E. coli* BL21(DE3) star and expressed overnight at 30°C in Terrific broth (TB). Cells were resuspended in 1X PBS and lysed by sonication. GST-TSG101_UEV_ was purified using glutathione affinity chromatography.

### Isothermal calorimetry

ITC experiments were performed in triplicate using a TA Instruments standard volume NanoITC in 20 mM sodium phosphate, pH 7.2, 150 mM NaCl, and 5% glycerol. Protein and ligands were buffer matched. ∼4 mM GRA14 PTAP (VQPTAPPAP) and HIV-1 Gag PTAP (PEPTAPPEE) synthetic peptides were injected in 2 μL aliquots to ∼94 μM protein. Heats of dilution were subtracted using buffer control titrations and data were fit to a one-site binding model and data was analyzed using NanoAnalyze software.

### ALG-2 and Pru::HA-GRA8 Immunoprecipitation

For ALG-2 co-IP, HFFs were cultured in 15-cm dishes and infected with Prustrain at a MOI of 3 for 24 h in three independent experiments. Dishes were washed three times with ice-cold PBS and monolayers were scraped from dishes in 1 mL of cold lysis buffer per 15-cm dish (50 mM Tris-HCl (pH 7.6), 200 mM NaCl, 1% Triton X-100, and 0.5% CHAPS), supplemented with cOmplete EDTA-free protease inhibitor cocktail (Roche Cat# 11836170001). Cellular lysates were passed through a 27G syringe 5 times and sonicated on ice. Lysates were incubated on ice for 30 min and spin down at 1,000 × g, 10 min, 4°C to remove debris. Proteintech PDCD6 rabbit polyclonal antibody (Cat # 12303-1-AP) was conjugated to Dynabeads M-270 epoxy (Invitrogen Cat# 14301) according to the manufacturer’s protocol. The supernatant (input) was incubated overnight at 4°C with 50µg of conjugated beads. For Pru::HA-GRA8 reciprocal co-IP, HFFs were infected with Pru::HA-GRA8 or untagged parental control (Pru) in parallel in triplicates at MOI of 3. The dishes post infection were processed as described above. The supernatants were incubated with 0.25 mg anti-HA magnetic beads (Pierce/Thermo Fisher, Cat# 88836; 100 µL slurry). Post incubation beads were separated on magnetic stand and washed four times in wash buffer (50 mM Tris pH 7.4, 300 mM NaCl, 0.1% Triton X-100). The target protein was then eluted with Laemmli buffer with 50 mM DTT, boiled for 5 min and separated on magnetic stand for 2 min. An aliquot of eluates was analyzed by immunoblotting. The protein for LC-MS/MS analysis was eluted using 0.1M Glycine pH 2.3 and the eluates were loaded, washed, and digested into peptides with 1 mg of trypsin on S-TRAP micro columns (ProtiFi, Farmingdale, NY) per manufacturer guidelines. S-TRAP peptide eluates were concentrated with a speed vac, desalted in HLB resin (Waters), and concentrated in a speed vac once more prior to LC-MS/MS acquisition.

### LC-MS/MS acquisition and analysis

Peptides samples derived from both ALG-2 and Pru::HA-GRA8 reciprocal co-IPs were resuspended in 10 µL of 0.1% trifluoroacetic acid (TFA) and loaded onto a Dionex RSLC Ultimate 3000 UHPLC system (Thermo Fisher Scientific, San Jose, CA) coupled online to an Orbitrap Fusion Lumos mass spectrometer (Thermo Fisher Scientific). The instrument was operated in data-dependent acquisition (DDA) mode. Full MS scans were acquired over an m/z range of 300–1,200 in the Orbitrap at a resolution of 120,000 (at m/z 200) with an AGC target of 5×10⁵. MS/MS spectra were acquired in the ion trap using top-speed mode (2 s cycle time), with an AGC target of 10⁴ and a higher-energy collisional dissociation (HCD) energy set to 30. Raw spectra were processed in Proteome Discoverer (v2.4, Thermo Fisher Scientific) using the SEQUEST search engine against the *T. gondii* ME49 proteome (ToxoDB release 59) and the Swiss-Prot human or mouse databases (January 2020). Searches included variable modifications of methionine oxidation and N-terminal acetylation and a fixed modification of carbamidomethyl cysteine. Trypsin was specified as the proteolytic enzyme, with precursor and product mass ion tolerances of 10 ppm and 0.2 Da, respectively. Peptide-spectrum match (PSM) and protein false discovery rates (FDRs) were set to 1%. For quantitative analysis, peptide intensity values were log2-transformed and normalized to the sample mean. Missing values were imputed from a normal distribution centered 2 standard deviations lower than the mean. Individual peptide fold changes (tagged versus control) calculated for a given protein and averaged to obtain protein fold enrichment. t-statistics were computed for each protein with at least two detected peptides by treating protein fold enrichment as the sample mean and using log-transformed peptide intensities to calculate the standard deviation, sample size, and degrees of freedom. Corresponding p-values were derived from t-distributions, and statistical significance across three biological replicates per condition was evaluated using two-tailed unequal variance t-test (p < 0.05) comparing peptide abundances between test and control samples. The proteomics raw files are available in the public repository ProteomeXchange (PRIDE) at the project number PXD078818.

### Super-resolution microscopy for TgGRA8, ALG-2, ALIX and TSG101

WT and HeLa cells expressing endogenously tagged GFP-TSG101 (36) were cultured on 1.5 mm coverslips in 6-well plates. To investigate the subcellular distribution of TgGRA8 in relation to ALG-2, ALIX, and TSG101, GFP-TSG101 HeLa cells were transiently transfected with pcDNA ALG-2-HA (28) and subsequently infected with ME49 Δ*ku80* parasites at a MOI of 5. TgGRA8, ALG-2, and TSG101 were detected using mAb anti-TgGRA8, anti-HA, and endogenous GFP tagging, respectively. To investigate the organization of TgGRA8 with ALG-2 and ALIX, WT HeLa cells were transfected with mCherry-ALG-2WT (52) and infected with Pru::HA-GRA8 parasites under similar conditions. Additionally, to examine the association of TgGRA8 with ALIX and TSG101, GFP-TSG101 HeLa cells were infected with Pru::HA-GRA8 parasites under similar conditions. TgGRA8 and ALIX were probed using anti-HA and purified monoclonal anti-ALIX antibody. Following infection, all the coverslips were fixed and stained for parasites using biotin-labeled polyclonal anti-*Toxoplasma* antibody respectively. Samples were imaged using a Nikon N-SIM super-resolution microscope equipped with a 100× objective.

### Multiple Sequence Alignment of GRA8 orthologs

The multiple sequence alignment (MSA) was performed for the TgGRA8 orthologs across tissue cyst forming Coccidian’s, *T. gondii* (TGME49_254720), *H. hammondi* (HHA_254720), *N. caninum* (NCLIV_008990), *B. besnoiti* (BbGRA8), and *C. suis* (CsGRA8) using MAFFT v7 with the L-INS-i algorithm for optimal accuracy (64). The resulting MSA was visualized and analyzed in Jalview v2.11 (65). Conservation and identity scores for each residue position in the ABEs were computed within Jalview. Residues were highlighted according to percent conservation and identity thresholds, with amino acid positions showing ≥30% conservation and ≥30% sequence identity across all aligned sequences marked as conserved. These thresholds were chosen to capture moderately conserved regions likely contributing to structural or functional roles while allowing visualization of sequence diversity. Alignments were colored using the ClustalX scheme to emphasize physicochemical similarities among residues.

### Membrane topology of TgGRA8 at the PVM

HFFs grown on circular micro cover glass were infected with Pru::HA-GRA8 strain at an MOI of 1 for 24 h. Infected cells were fixed with 4% PFA for 10 min and washed one time with PBS and PFA quenching performed using 0.1M glycine at RT for 5 min and blocked for 20 min in 3% BSA. After one wash the cells were permeabilized with saponin 0.01% for 10 min and were then stained for 1 h with primary rabbit anti-TgSAG1 and rat anti-HA (clone 3F10, Sigma-Aldrich) antibodies following secondary Ab staining. Coverslips were imaged at 63x using a Leica SP5 confocal microscope. PVs were located using differential interference contrast microscopy.

### ALG-2 expression and purification

A truncated human ALG-2 construct lacking the N-terminal Gly/Pro-rich region (previously described as des3-23; (37) was generated for bacterial expression. The des3-23ALG-2 sequence was cloned in MCS of the pETite-NHIS vector expressing 6XHis tag at the N-terminal. Site-directed mutagenesis was performed to generate the des3-23ALG-2^ΔGF122^(ALG-2^ΔGF122^) variant using the Q5 Site-Directed Mutagenesis Kit (NEB #E0554S) and primers F: GCTACCGGCTCTCTG and R: CTGAGAGGGCCTGCTTC. Recombinant proteins were expressed in *E. coli* HI-Control BL21(DE3) cells at 25°C for 18-20 h. Following expression, proteins were purified by Ni²⁺-affinity chromatography using an AKTA Pure FPLC system and dialyzed with 20 mM Tris and 20 mM NaCl. Protein purity was evaluated by SDS-PAGE and quantification was performed using Bicinchoninic acid assay (BCA).

### Bio-layer Interferometry

For biolayer interferometry (BLI) experiments, TgGRA8 (GRA8-ABE1: Biotin-PEG2-VRPPPPGFHPSVIPNPPYPLGTPAGMPQPEVPPL; GRA8-Poc1,2: Biotin-PEG2-IPNPPYPLGTPAGMPQPEV; GRA8-Poc3: Biotin-PEG2-VRPPPPGFHPSV) and ALIX (Biotin-PEG2-QGPPYPTYPGYPGYSQ) peptides were synthesized with an N-terminal biotin and a PEG2 linker (Peptide 2.0 Inc.). Peptide identity and purity were verified by HPLC and mass spectrometry. Lyophilized peptides were resuspended in molecular-grade water and stored at - 20°C until use. BLI measurements were performed on an Octet Red instrument (Sartorius) maintained at 25°C with orbital shaking at 1,000 rpm. The assay buffer consisted of 20 mM Tris, 20 mM NaCl, and 0.05% Tween-20, supplemented with either 100 µM CaCl₂ or 1 mM EGTA. Streptavidin (SA) biosensor tips (Sartorius #18-5019) were pre-equilibrated in assay buffer for 10 min prior to peptide loading. Assays were conducted in 96-well plates, each well containing 200 µL of either assay buffer, biotinylated peptide, or purified protein diluted in assay buffer. Serial dilutions of the studied protein were prepared immediately before the assay. For ALIX peptide measurements, des3-23 ALG-2 was tested over a concentration range of 3.12–0.39 µM. For assays using the TgGRA8 and ALIX peptide, rALG-2^WT^ and rALG-2 ^ΔGF122^ dilutions ranged from 73 nM to 12 nM. The standard experimental workflow consisted of sequential steps: baseline (60 s), peptide loading (1200 s), wash (900 s), secondary baseline (60 s), association (300 s), and dissociation (2000 s). Binding kinetics and affinities (K_d_) were obtained by global fitting to a 1:1 binding model using the Octet Red data analysis software.

### Size exclusion chromatography

Affinity-purified recombinant rALG-2^WT^ was analyzed alone or after mixing with synthetic TgGRA8-ABE1 peptide at a 1:1 equimolar ratio under reducing conditions in the presence of 5mM β-mercaptoethanol. Samples were loaded onto a Superdex 200 Increase 10/300 GL size-exclusion chromatography column equilibrated in 20 mM Tris and 20 mM NaCl and eluted in the same buffer at a flow rate of 0.5 mL/min. Low-molecular-weight standards and blue dextran were used to estimate apparent molecular mass and column void volume, respectively. For the ALG-2-TgGRA8-ABE1 complex mixture, buffer elution was performed first to assess soluble complex recovery, followed by 0.5N NaOH flushing to recover bound peptide-protein complex on the column.

### AlphaFold modelling of TgGRA8 ABE-I and ALG-2

The TgGRA8 ABE-I (VRPPPPGFHPSVIPNPPYPLGTPAGMPQPE VPPL; TGME49_254720) and ALG-2 (MAAYSYRPGPGAGPGPAAGAALPDQSFLWNVFQRVDKDRSGVISDTELQQALSNGTWTPFNPVTVRSIISMFDRENKAGVNFSEFTGVWKYITDWQNVFRTYDRDNSGMIDKNELKQALSGFGYRLSDQFHDILIRKFDRQGRGQIAFDDFIQGCIVLQRLTDIFRRYDTDQDGWIQVSYEQYLSMVFS IV) structure prediction was performed using AlphaFold3 (66). Structural visualization of ALG-2-binding motifs and predicted binding interfaces were performed in UCSF ChimeraX https://www.rbvi.ucsf.edu/chimera (67).

### Generation of mCherry ALG-2 pocket mutants

Plasmids encoding mCherry-ALG-2^WT^, mCherry-ALG-2^ΔGF122^, and mCherry-ALG2^E47A/E114A^ were described previously (52). Additional point mutations were introduced into the mCherry-ALG-2 ^WT^, and mCherry-ALG-2^ΔGF122^ constructs to generate mCherry-ALG-2^F85A^, and mCherry-ALG-2^ΔGF122 F85A^ mutants. Mutagenesis was performed using the Q5 Site-Directed Mutagenesis Kit (NEB #E0554S). For a full list of primers used for generation of mCherry ALG-2 pocket mutants, please refer to **Table 1**

### Targeted metabolomics and comparative analysis

HFFs were infected with equal numbers of RH*Δku80*WT, RH*Δku80Δgra8*, and RH*Δku80Δgra8GRA8* parasites and harvested at 44 h post-infection. Parasites were harvested and analyzed by targeted bulk metabolomics using LC-MS/MS as described previously (68). Detected metabolites were normalized prior to downstream analysis, and the resulting dataset was compared with previously published RH*Δgra14* and RH*Δcpl* metabolomics datasets (32). PCA was performed using normalized metabolite abundance values to assess global metabolic differences among different strains. Heatmaps were generated to visualize relative metabolite changes across strains; metabolite abundances in mutant strains were expressed as log2 fold change relative to RH*Δku80*WT. Pathway enrichment analysis of metabolites altered in RH*Δku80Δgra8*, RH*Δgra14* and RH*Δcpl* parasites relative to RHΔku80WT was performed using MetaboAnalyst (69). Enriched pathways are shown as per the enrichment factor.

## Abbreviations

ESCRT: endosomal sorting complex required for transport
ABE: ALG-2-binding element
ABM: ALG-2-binding motif
ALG-2: apoptosis-linked gene 2
ALIX: ALG-2-interacting protein X
TSG101: Tumor susceptibility gene 101
Tg: *Toxoplasma gondii*
PV: parasitophorous vacuole
PVM: parasitophorous vacuole membrane
GRA: dense granule protein
BLI: Bio-layer Interferometry
SEC: Size exclusion chromatography
LC-MS: Liquid chromatography-mass spectrometry

## Acknowledgments

We thank Dr. John Boothroyd for providing pcDNA-ALG-2-HA plasmid, Dr. Gary Ward for providing the TgGRA8 antibody, Dr. Peter Bradley for providing the RHΔΔΔ*gra14* strain, and Dr. Schuyler B. van Engelenburg for sharing the endogenously tagged GFP-TSG101 HeLa cells. We also thank Dr. Nicole Koropatkin for access to an ITC instrument. We thank Dr. Simone Sidoli and the Einstein Laboratory for Macromolecular Analysis and Proteomics for all their assistance with LC-MS/MS preparations and analysis. We also thank Isabelle Coppens and Julia Romano for helpful suggestions and Tracey Schultz for laboratory support.

## Disclaimer

This work was prepared while Alfredo J. Guerra was employed at the University of Michigan. The opinions expressed in this article are the authors’ own and do not reflect the view of the National Institutes of Health, the Department of Health and Human Services, or the United States government.

## Supporting information Figure legends

### Supporting information

**S1 Fig. Loss of TgGRA14 or TgGRA64 has minimal effect on ALG-2, ALIX, and TSG101 recruitment to the PVM.** ALG-2-HA transfected, WT, and GFP-TSG101-expressing HeLa cells were infected with either RH (WT) or RHΔ*gra14* **(A)** and RHΔ*ku80*WT, RHΔ*ku80*Δ*gra64*, or RH*Δku80*Δ*gra64*Δ*gra14* parasites **(B)**. ALG-2 and ALIX were detected using anti-HA and anti-ALIX antibodies, respectively and while TSG101 was endogenously tagged with GFP. Each data point represents a single well intensity ± SEM. Statistical analysis was performed using unpaired two-tailed t-tests **(A)** or one-way ANOVA with multiple comparisons **(B)** as indicated above each plot.

**S2 Fig. Loss of TgGRA8 markedly reduced ALG-2 and ALIX recruitment, with minimal effects on TSG101 across replicates and parasite strains. (A-C)** Representative confocal micrographs of ME49, ME49Δ*gra8*, and ME49Δ*gra8*::*GRA8* parasites showing recruitment of ALG-2, ALIX, and TSG101 on the PVM. Host cell nuclei were stained with DAPI, and parasites were visualized using anti-*Toxoplasma* antibody. ALG-2, ALIX and TSG101 were detected using similar staining strategy as applied for RHΔku80WT parasites. Scale bar, 20 µm. **(D-F)** Quantification of ALG-2 (D), ALIX (E), and TSG101 (F) recruitment represented as parasite-to-cytoplasm mean fluorescence intensity ratios. Data represents SEM from 3 independent biological replicates (>10,000 PVs each; ∼16 wells/replicate). Each point represents the well average (∼16 wells per biological replicates) ± SEM. Statistical comparisons were performed using one-way ANOVA with Tukey’s multiple comparisons test.; p-values are shown above the bars.

**S3 Fig. ALG-2, ALIX, and TSG101 recruitment across individual wells and biological replicates analyzed in ME49 and RH parasites.** ALG-2-HA transfected, WT, and GFP-TSG101-expressing HeLa cells were infected with RHΔ*ku80*WT, RHΔ*ku80*Δ*gra8*, or RHΔ*ku80*Δ*gra8*::*GRA8* (A) and ME49, ME49Δ*gra8* or ME49Δ*ku80*Δ*gra8*::*GRA8* parasites **(B)**. ALG-2 and ALIX were detected using anti-HA and anti-ALIX antibodies, respectively and while TSG101 was endogenously tagged with GFP. Each data point represents a single well intensity ± SEM. Statistical analysis was performed using one-way ANOVA with multiple comparisons as indicated above each plot.

**S4 Fig. GRA8-dependent recruitment of endogenous ALG-2 to the PV.** Representative IFA images of HeLa cells infected with RHΔ*ku80*WT, RHΔ*ku80*Δ*gra8*, or RHΔ*ku80*Δ*gra8*::*GRA8* **(A)** and ME49, ME49Δ*gra8* or ME49Δ*ku80*Δ*gra8*::*GRA8* parasites. Scale bar, 5 µm. **(B)**. Endogenous ALG-2 was detected using ALG-2 Polyclonal Ab. Zoomed insets show ALG-2 enrichment at the PV in WT and Δ*gra8*::GRA8 parasites in both type I and II, with reduced recruitment in Δgra8 parasites. Scale bar, 2 µm.

**S5 Fig. GFP-TSG101 expressing HeLa cells were infected with Pru::HA-GRA8 parasites and stained for TgGRA8 (blue), and ALIX (red).** SIM images showing the spatial proximity of TgGRA8, ALIX, and TSG101 (green) at the PVM. Merged images and magnified insets highlight discrete regions of colocalization (arrows) between ALIX and TSG101, as well as sites where all three proteins overlap. Scale bar, 2 µm.

**S6 Fig. TgGRA8 ABE-1 peptides binds to rALG-2** ^ΔGF122^ **in Ca^2+^-dependent manner.** BLI showing the binding of des3-23ALG-2 ^ΔGF122^ to N-terminally biotinylated ALIX **(A)**, GRA8-Poc1,2 **(B)**, GRA8-Poc3 (**C)**, GRA8-FL **(D)** peptides under either Ca²⁺-replete (100 μM CaCl₂) or Ca²⁺-chelated **(D)** peptides under either Ca²⁺-replete (100 μM CaCl₂) or Ca²⁺-chelated (1 mM EGTA) conditions. The analysis involved exposing these peptides to an array of different concentrations of des3-23ALG-2 ^ΔGF122^ (concentrations shown in the key). The association between the peptides and proteins for both the experiments were measured for 300 s followed by dissociation for 2,000 s. Data shown here are from two independent experiments conducted for each condition. BLI sensorsgrams were baseline-corrected and globally fitted using a 1:1 binding model.

**S7 Fig. mCherry-ALG-2^WT^ or mCherry-ALG-2^F85A^ or mCherry-ALG-2^ΔGF^ or mCherry-ALG-2^ΔGF/F85A^ or mCherry-ALG2^E47A/E114A^ recruitment across individual wells and biological replicates analyzed in RH and ME49 parasites.** HeLa cells were transfected with mCherry-ALG-2^WT^ or mCherry-ALG-2^F85A^ or mCherry-ALG-2^ΔGF^ or mCherry-ALG-2^ΔGF/F85A^ or mCherry-ALG2^E47A/E114A^ and infected with either RHΔku80WT **(A)** or ME49Δku80WT **(B)** parasites to study the recruitment of ALG-2. Each data point represents a single well intensity ± SEM. Statistical analysis was performed using one-way ANOVA with multiple comparisons as indicated above each plot.

**S8 Fig. Loss of GRA8 alters *T. gondii* metabolic pathways. (A)** Dot plot showing mean log2 fold change in metabolite abundance relative to WT parasites for Δ*cpl*, Δ*gra1*, Δ*gra8* and Δ*gra8GRA8* parasites relative to WT. Each point represents an individual metabolite; filled circles denote metabolites significantly altered compared with WT, whereas open circles indicate non-significant changes. **(B, C)** Pathway-enrichment analysis of metabolites altered relative to RHΔ*ku8* WT was performed using MetaboAnalyst (69) for **(B)** RHΔ*gra14* and **(C)** RHΔ*cpl*. Enriched pathways are plotted by enrichment factor; point color indicates significance as −log10(FDR), and point size indicates pathway impact.

**S1 Table. Primers used in this article.** The table consist of list of primers, sgRNAs and donor oligonucleotides used for generating GRA8 knockout, endogenous HA tagging, genetic complementation and validation of genome editing in *T. gondii*, as well as primers used to generate ALG-2 mutant constructs. Sequences are shown in the 5′-3′ orientation.

**S1 Data. LC-MS/MS data from ALG-2 Co-IP experiments.** The data lists the calculated protein fold change (Pru/Control) and log2 P values for all proteins detected in each replicate. The proteomics raw files are available in the public repository ProteomeXchange (PRIDE) at the project number PXD078818.

**S2 Data. LC-MS/MS data from Pru::HA-GRA8 Co-IP experiments.** The data lists the calculated protein fold change (Pru::HA-GRA8/Control) and log2 P values for all proteins detected in each replicate. The proteomics raw files are available in the public repository ProteomeXchange (PRIDE) at the project number PXD078818.

**S3 Data. Targeted metabolomics data from GRA8 parasite strains.** Raw metabolite abundance values measured by LC-MS/MS for WT, Δgra8, and Δgra8*GRA8* parasites. Each column represents an independent biological replicate, and each row corresponds to an individual metabolite detected in the targeted metabolomics panel.

